# Neuroelectrophysiological correlates of self-reported extended cessation of consciousness in advanced meditators: A multimodal EEG and MEG study case series

**DOI:** 10.1101/2025.09.19.677455

**Authors:** Kenneth Shinozuka, Winson F.Z. Yang, Ruby M. Potash, Terje Sparby, Matthew D. Sacchet

## Abstract

In some contemplative traditions, “extended cessation” (EC) refers to a state of advanced meditation in which the meditator self-reports intentionally suppressing their own consciousness and re-emerging with a profound sense of clarity and equanimity. Here, we present the first electrophysiological study of EC, in which five meditators underwent concurrent electroencephalography (EEG) and magnetoencephalography (MEG) recording. EC significantly increased high-frequency global coherence in all five of the participants, yet it did not consistently affect Lempel-Ziv complexity or power across participants. Our results provide preliminary evidence that the neural signatures of EC may be distinct from other states that are characterized by a loss of consciousness, including deep sleep and anesthesia. EC may not be simply “switching off” consciousness but rather reorganizing the neural dynamics that underlie consciousness.

## Introduction

Using novel techniques to classify and correlate phenomenology with neural markers, studies in the recent “third wave” of research on meditation have found that advanced meditators can access altered states of consciousness characterized by profound clarity, peace, and equanimity (*1–8*). Advanced meditation research also seeks to explain the profound shifts in consciousness that can arise from contemplative mastery. One such shift, which we call extended cessation (EC), refers to a state reported as rare and temporary suspension of ordinary consciousness (*9*, *10*). That is, after specific and especially deep meditation, all perception, feelings, and thoughts – three mental factors regarded as universally present in conscious cognition (*11–14*) – cease entirely for an extended period, which can range from minutes to hours. In Theravada Buddhism, *nirodha samāpatti* is a form of EC that represents a self-reported cessation of all mental and sensory activity, and is considered to be among the highest meditative attainments, or the highest meditative attainment (*12*, *15*, *16*). However, due to ongoing debate regarding the criteria for and implications of nirodha samāpatti, we adopt a tradition-neutral term, extended cessation (EC), to refer to the self-reported, intentional, and sustained non-conscious states reported here (*17*, *18*). EC is phenomenologically distinct from deep sleep, anesthesia, and disorders of consciousness (e.g., coma) for several reasons: EC is intentional; there is no sense of time passing during EC; one cannot be awakened by external stimuli; and meditators typically report experiencing feelings of immense clarity and peace after coming out of EC (*12*, *13*). These features suggest that EC constitutes a unique, volitionally induced state that may serve as a novel model system for studying the boundaries and mechanisms of conscious awareness. Preliminary evidence indicates that momentary cessations, which last on the order of seconds, are associated with pre- and post-cessation decreases in global alpha power and functional connectivity while elevating whole-brain criticality (*9*, *17*, *19*). However, these changes in the brain were observed before and after the cessation, rather than during the cessation itself, which no research has characterized to date.

Thus, a central question is whether the neural correlates of EC (volitionally induced loss of consciousness) differ from those of deep sleep, anesthesia, and disorders of consciousness (unintentional loss of consciousness). To address this, we examined three electrophysiological biomarkers of consciousness, both globally and regionally in the brain: spectral power, complexity (Lempel-Ziv complexity [LZc] and integrated information), and global coherence.

These markers have well-characterized profiles in unconscious states. For example, loss of consciousness tends to amplify power in lower frequencies (delta, theta bands) while suppressing power in higher frequencies (beta, gamma bands) (*20–23*). Additionally, loss of consciousness reduces the complexity of neural activity, as measured by LZc (*24–27*). Finally, anesthesia decreases alpha-band global coherence, which measures how much coherence can be explained by the most dominant phase-locked network at that frequency (*28*).

What is the neuroelectrophysiology of EC, and does EC affect the brain differently than deep sleep, anesthesia, and disorders of consciousness? In this first electrophysiological study of EC, conducted in parallel with a functional neuroimaging study on the same participants (*29*), we compared brain activity during EC to brain activity during two control tasks in five advanced meditators. We hypothesized that, like other states in which consciousness is absent, EC would decrease complexity and alpha-band global coherence. Based on previous case reports of brief cessations, we also hypothesized that EC would decrease alpha power.

## Methods

### Participants

This study included five advanced meditators, hereafter referred to as sub001 through sub005. Sub001 (male [M]; 52 years old) had over 25 years of meditation experience with an estimated total lifetime meditation practice of 20,000+ hours at the time of data collection. Sub002 (M; 32 years old) had over 19 years of meditation experience and, at the time of data collection, an estimated total lifetime meditation practice of 15,000+ hours. Sub003 (M; 46 years old) had over 4 years of meditation experience and an estimated total lifetime practice of 10,000+ hours at the time of data collection. Sub004 (M; 66 years old) had over 50 years of meditation experience and an estimated total lifetime practice of 25,000+ hours at the time of data collection. Finally, Sub005 (M; 66 years old) had over 40 years of meditation experience and an estimated total lifetime practice of 25,000+ hours at the time of data collection. We note that lifetime hours are approximate and do not necessarily correlate to meditative expertise (*30*). Important to the current study, the participants are advanced meditators and reported being able to reliably practice EC. The study was approved by the Mass General Brigham Institutional Review Board (IRB), and all participants provided informed consent.

### Study design

#### Extended cessation

EC data collection runs were acquired within a broader study of advanced meditation. Participants were asked to enter EC with the intention of maintaining the state for approximately 15 min. Participants completed a varying number of runs depending on ability and fatigue (see **Table S1**). The actual duration of each EC run lasted anywhere from 1 minute to 36 minutes, depending on the participant. Note that all participants prepared by meditating in the EEG/MEG, for varying amounts of time, before they reached EC. Timestamping of entry into and exit out of the EC state are described in *Section 2.4*. Participants also completed a standardized EC phenomenology questionnaire designed to capture the typical phenomenology of their EC practice. The questionnaire asked participants to report how they generally prepare for, enter into, experience, and exit from EC, as well as any aftereffects they commonly observe. For each phase, participants responded “yes” or “no” to a series of items reflecting experiences they personally associate with EC and were not limited to the experimental session. Three participants provided their phenomenology responses after data collection had formally concluded, and their delayed reports have been included in the dataset.

#### Non-meditative control conditions

Two non-meditative control conditions were included to engage the participants non-meditative cognitive processes. These controls were designed to avoid the induction of meditative states. Resting-state controls were not used due to concerns that advanced meditators may enter meditative states at rest (*31–37*). Non-meditative controls included two tasks: a Counting task, in which the participant was asked to sublingually count down (i.e., mentally without moving the lips) in decrements of five from the number 10,000 for eight minutes; and a Memory task, in which the participant was the participant was asked to reminisce the events of the past two weeks and narrate them in their mind for eight minutes. Eyes were closed during both non-meditative conditions as well as during EC. Each participant completed two runs of each control task, except for sub002, who completed one run of each.

As noted below in ***Statistical analyses***, control conditions were pooled together in order to avoid multiple comparisons; however, comparisons with each control condition separately were always reported in the **Supplementary Materials**.

#### Data acquisition

EEG data was acquired from one participant (sub001), and combined MEG-EEG data was acquired from the other four participants (sub002-sub005). For sub004, a substantial portion of EEG electrodes showed flat signals compared to the remaining electrodes. As a result, their EEG data was excluded from analysis and only the MEG data was analyzed. For sub001, continuous EEG activity was recorded from a customized 96-channel actiCAP system using an actiCHamp amplifier (Brain Products GmbH, Gilching, Germany). Impedances were kept below 5 kΩ. The ground (GND) channel was embedded in the cap and was located anterior and to the right of Channel 10, which roughly corresponds to electrode Fz. Channel 1 (Cz) served as the online reference channel during data acquisition. All signals were digitized at 500 Hz using BrainVision Recorder software (Brain Products). Continuous MEG-EEG activity was recorded on the TRIUX system at Massachusetts General Hospital with 306 MEG channels (102 Magnetometers, 204 Gradiometers), 2 EOG, and a varying number of simultaneously collected EEG channels. For sub002, 70 EEG channels were recorded, whereas for the other three participants, EEG activity was recorded from a customized 128-channel MEG BrainCap system. The MEG-EEG system was always located in an Imedco magnetically shielded room, with a shielding factor of approximately 250,000 at 1Hz. Sampling frequency was 1000 Hz with a high bandpass of 0.1 Hz and a low bandpass of 330 Hz.

Note that fMRI data, which are reported in another study (*29*), were acquired from three of the participants in separate sessions than EEG and/or MEG.

#### Preprocessing

Both EEG and MEG data from the EC condition were firstly truncated to the period of time in which the participants were in EC, rather than meditating in preparation to enter EC. Given the absence of consciousness during EC, real-time indication of entry was not possible for all participants. Indications of exit, typically through a delayed motor response, varied across individuals. Sub002 used the button press to indicate entry and exit out of EC; sub004 pressed a button to indicate entry into EC (the recording ended about 30 seconds after exit from EC); and sub005 pressed a button to indicate exit from EC (EC started 1-2 minutes beforehand, depending on the run). Sub001 and sub003 did not press buttons. Instead, the timings of sub001’s entry into and exit out of EC were inferred based on the subject’s report of their eyeblinks prior to the EC onset (which were determined by visible artifacts in the EEG data). Sub003 reported that EC began approximately one minute into the recording, which was stopped 30 seconds after the participant signaled that they had exited EC.

EEG and MEG data were preprocessed using the FieldTrip software (*38*). Both EEG and MEG data were high-pass filtered at 1 Hz, downsampled to 200 Hz, and line noise at 36, 60, and 72 Hz was removed with a notch filter. High-amplitude outliers (*z* > 20) were removed using partial rejection and a jump-preserving median filter (order 9). Bad channels were rejected by visual inspection. Independent component analysis (Infomax algorithm) was applied separately to the EEG and MEG data (*39*); then, components corresponding to heartbeat, muscle, and eye blink artifacts were identified and removed based on their scalp topography and time course.

#### Source reconstruction

Source analysis was performed with exact low-resolution electromagnetic tomography (eLORETA), as implemented in FieldTrip (*40*). For each participant, preprocessed sensor data from all runs were restricted to the channels common to every run of that participant (sub001: 90 EEG; sub002: 69 EEG + 306 MEG; sub003: 117 EEG + 303 MEG; sub004: 306 MEG; sub005: 122 EEG + 298 MEG). EEG data were re-referenced to the common average of this final channel set. Each participant’s anatomical MRI was coregistered to the sensor coordinate system (initial fiducial-based alignment refined with digitized head-shape points), and the source model was defined as the MNI centroids of the 100-parcel Schaefer atlas, inversely warped to the individual anatomy (*41*). Volume conduction was modeled with a concentric-spheres model for EEG and a corrected single-shell model for MEG (*42*). Leadfields were computed with column-wise normalization (order 1) and, for MEG, rank reduction to two; for participants with combined EEG-MEG, the EEG data were scaled by the ratio of the mean channel-wise standard deviation of the MEG to that of the EEG (computed over all runs), and the EEG and MEG leadfields were concatenated per source position.

eLORETA spatial filters were computed from the concatenated leadfields with a regularization parameter of 0.05. As eLORETA yields a three-dimensional dipole moment per source, a fixed orientation was assigned to each source as the dominant eigenvector of the filter-projected sensor covariance. Applying the resulting 100 spatial filters to each full-length run yielded one parcel time series per Schaefer region at 200 Hz, which formed the basis of all subsequent analyses.

#### Global coherence

The cross-spectral density matrix (CSPD) was computed using a Hamming window of 1 second, 50% overlap between windows, and the number of discrete Fourier transform points equal to the amount of between-window overlap. At each frequency from 1 to 50 Hz, global coherence was computed as the leading eigenvalue of the CSPD at that frequency divided by the sum of all the eigenvalues (*28*). The leading eigenvector of the CPSD at each frequency was also determined. We reported changes in the leading eigenvector at 11 Hz, following previous literature (*28*).

Because increases in beta- and gamma-band global coherence could in principle reflect cranial muscle (EMG) activity rather than neural synchronization, we performed two control analyses. First, a spectral-extension test exploited the fact that tonic cranial EMG is spectrally broadband, with power extending well beyond 60 Hz, whereas neural gamma oscillations are band-limited [refs A–C]. For each participant we computed, on the raw sensor-space recordings, the per-channel log-ratio of cessation to control power in an analysis band reflecting neural activity (39–50 Hz, matching the frequency range of the significant gamma-coherence clusters) and in an EMG-index band (62–95 Hz, excluding the 71–73 Hz line-noise band). Two hypotheses were tested per participant: (a) whether 62–95 Hz power differed between cessation and the pooled control conditions and (b) whether the cessation-related change in 39–50 Hz log-power exceeded that in 62–95 Hz log-power.

Second, we tested whether the parcel-wise spatial distribution of the gamma-band effect followed an EMG-like topography, computing Spearman correlations between each parcel’s effect size and (i) its proximity to the scalp sites of maximal cranial-muscle contamination, defined as the minimum distance from each parcel’s MNI centroid to the standard 10-20 positions of electrodes Fp1/Fp2 (overlying the frontalis) and F7/F8/T7/T8 (overlying the temporalis), the sites at which EMG contamination is maximal (*43*); and (ii) its radial distance from the head center. Because parcel-level maps are spatially autocorrelated, significance was assessed with a volumetric spin permutation (*44*, *45*).

#### Power spectra

Source-space power spectral densities (PSDs) were computed based on Welch’s overlapped segment averaging estimator, with windows of 200 samples in length, 50% overlaps between windows, and the number of discrete Fourier transform points used for the PSD estimates equal to the amount of between-window overlap. We measured the spatial distribution of power in the alpha band (8-13 Hz), due to prior literature linking regional alpha to absence of consciousness.

### Complexity

#### Lempel-Ziv complexity

Each region’s timeseries was binarized, such that timepoints above the mean were classified as “1” and timepoints below were classified as “0”. Lempel-Ziv complexity (LZc) was computed on the binary timeseries using the 1976 algorithm (*46*). LZc was then normalized by the length of each data segment, since LZc tends to be higher for longer segments. In particular, we multiplied LZc by log_2_(T)/T, where T is the length of the segment (*47*).

Changes in LZc sometimes arise from alterations in the power spectrum rather than the dynamics of the timeseries. Therefore, previous work has normalized LZc by the LZc of a phase-shuffled timeseries, in which the power spectrum is preserved but the dynamics are completely scrambled (*48*). We refer to LZc divided by the LZc of a phase-shuffled timeseries as “phase-normalized LZc.”

Both LZc and phase-normalized LZc were measured globally by averaging across regions. Regional estimates of LZc are also reported. Only broadband LZc and phase-normalized LZc were computed.

#### Integrated information

Integrated information theory claims that consciousness can be quantified as Φ, which, broadly speaking, captures the amount of information that is lost when a system is partitioned into subsystems. In other words, Φ is the extent to which the information in the whole system is greater than that of its parts. Φ is difficult to measure due to the large number of ways of partitioning complex systems. To address this challenge, Barrett and Seth proposed an empirical Φ measure that is more computationally tractable (*49*):

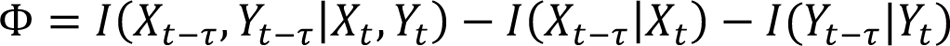

For a system with two components *X* and *Y*, empirical Φ is the amount of information that the pasts of *X* and *Y* jointly encode about their present values, above and beyond the amount of information that the pasts of *X* and *Y* separately encode about their present values.

We measured empirical Φ for each pair of regional timeseries and then averaged across all pairs. τ was set to 1 sample.

### Statistical analyses

Due to the small number of participants in this study (*N* = 5), we performed subject-level analyses rather than group-level analyses, and then we qualitatively analyzed trends across subjects, as done in extensive prior research with small samples, including humans and non-human primates (*28*, *51–54*). Each measure was computed in non-overlapping 15-second epochs of the source-reconstructed data for all participants. Epochs whose peak-to-peak amplitude or variance (averaged across parcels) deviated from the within-run median by more than five standard deviations were excluded (**Supplementary Table 1** reports epochs analyzed per participant and condition; because sub005’s EC runs were short, only 7 total EC epochs were available for that participant, and its results should be read with correspondingly reduced power in mind.)

Because consecutive epochs from a continuous run are not statistically independent (the lag-1 autocorrelation of epoch-level band power reached *r* = 0.74 in some runs), condition comparisons used block permutation rather than epoch-level permutation. That is, contiguous 60-second blocks of epochs within each run served as the exchangeable units, epoch condition-labels were permuted (5,000 permutations), and the test statistic was the difference of condition means (two-sided), weighted by the sum of block lengths. Results were unchanged across block lengths of 30–120 s.

Our primary comparison contrasted the EC condition against the two control conditions pooled. However, a Counting-versus-Memory equivalence check (same block-permutation procedure) was run per participant and measure, and no matter what, the separate per-control comparisons are reported in the **Supplementary Materials**.

For band-averaged power, we compared mean power within the five canonical frequency bands (delta 1–4, theta 4–8, alpha 8–13, beta 13–30, gamma 30–48 Hz) and corrected across bands with the False Discovery Rate (FDR). As a supplementary analysis, we also corrected power across individual frequencies ranging from 1–50 Hz. Full global-coherence spectra (1–50 Hz, 1-Hz resolution) were assessed with cluster-based permutation tests (*53*). Global-coherence values within 33–38 Hz were excluded from inference, as the 35–37 Hz band-stop filter renders coherence estimates in this range artifactual. The spatial distribution of alpha power was corrected across the 100 parcels with FDR.

## Results

### Phenomenology of extended cessations

A table reporting the phenomenology of different stages of EC is shown in **Figure S1.** All participants used advanced concentration absorption meditation *jhanas* (ACAM-J) to prepare for EC. ACAM-J is a series of concentration states that includes strong and stable focus, blissful sensations, emotional happiness, and peacefulness, followed by a progressive reduction of the contents of consciousness. With one exception, participants involved elements of advanced investigative insight meditation (AIIM), that is, a close, neutral observation of phenomena, as part of the setup of EC (*37*).

All participants reported that internal verbalization (thinking), body sensations, and feeling ceased during the onset of EC. During EC, conscious experience was completely absent. When exiting EC, all participants reported that autobiographical memory and ego-centered cognitive processing was absent, and there was no awareness of a separation between oneself and the environment. With one exception, all participants reported a strong clarity of mind when exiting EC, the presence of a pristine awareness without an object, and the re-emergence of mind, bodily sensation, and internal verbalization (in that order). After EC, all participants experienced extreme sensory vividness, but also a strong and lasting peace. They all also reported being inclined to solitude. With one exception, an experience of deep joy was also reported.

While the core phenomenological features of EC were consistent across individuals, participants noted that the depth or clarity of the experience may have been attenuated during scanning, possibly due to the nature of the recording environment (e.g., discomfort arising from wearing an EEG cap, etc.).

### EC significantly increases high-frequency global coherence in all participants

EC significantly elevated global coherence at beta- and/or gamma-band frequencies in all five participants (**Figure 2**; statistics are displayed in **Table 1**). The exact frequency range varied based on the participant. Global coherence was significantly higher on EC at 12–31 Hz and 45–50 Hz for sub001; 39–45 and 47–50 Hz for sub002; 23–32 Hz and 39–50 Hz for sub003; 30–32 Hz and 39–50 Hz for sub004; and 39–44 Hz for sub005. Global coherence was significantly different between control conditions for sub002 at several frequencies; the per-control analysis revealed that global coherence on EC was significantly higher compared to Counting at 30–32 Hz and 39–50 Hz, but not compared to Memory (**Figure S2**).

**Figure 1.**
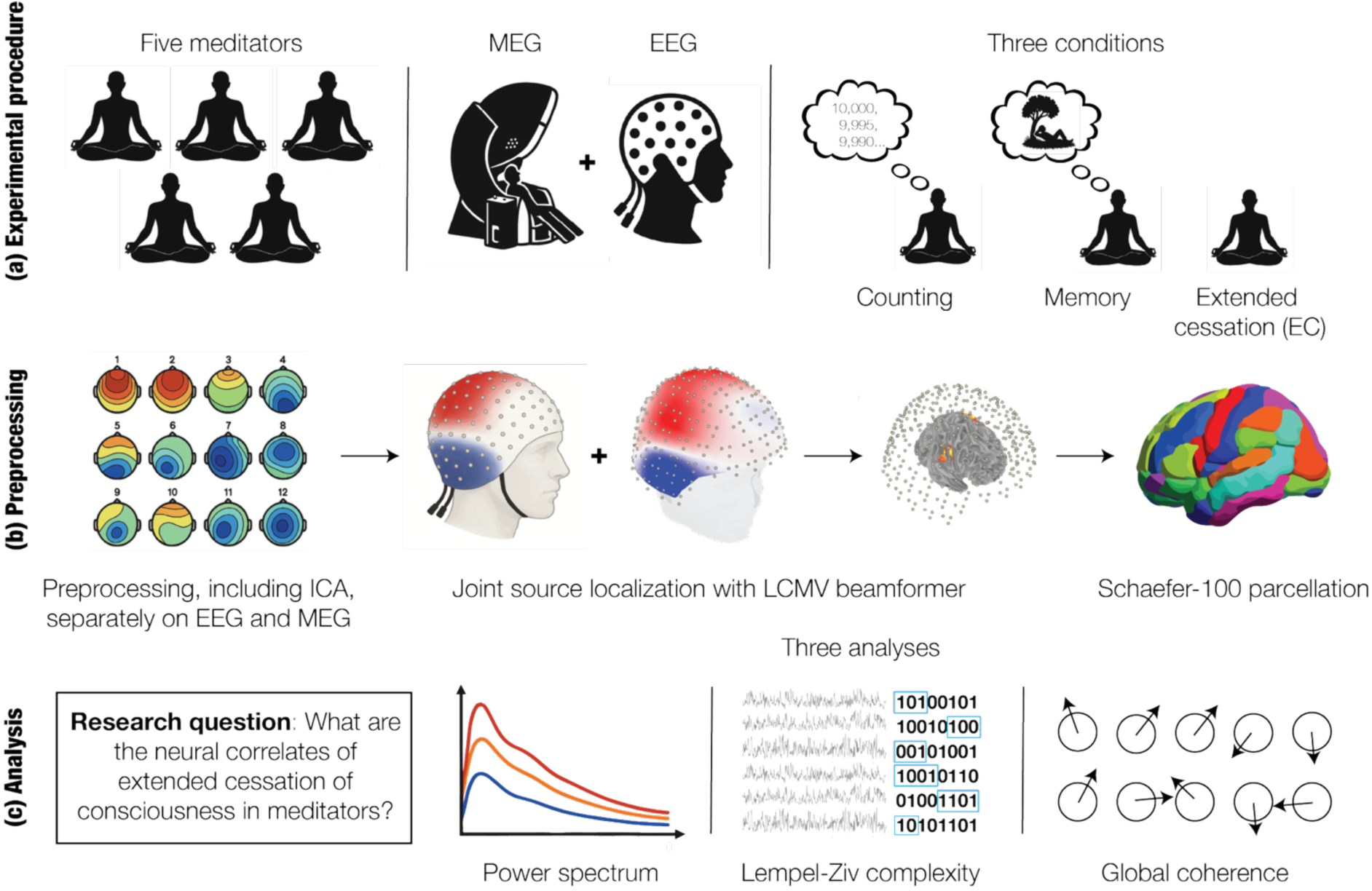
Overview. (a) Five advanced meditators were examined in three different conditions: extended cessation of consciousness (EC), a control Memory task in which participants reminisced on recent events, and a control Counting task in which participants counted down from 10,000. One participant was scanned with just EEG; for another participant, only MEG data was analyzed; and for the remaining three participants, simultaneous EEG and MEG data were acquired and analyzed. (b) EEG and MEG data were preprocessed separately, and then joint source reconstruction was performed by concatenating the respective leadfields. The sources were the centroids of the regions in the Schaefer-100 parcellation. (c) All analyses were conducted in source-space. Power spectra, Lempel-Ziv complexity, and global coherence were measured. These measures were chosen because they have been previously analyzed in research on loss of consciousness.

**Figure 2.**
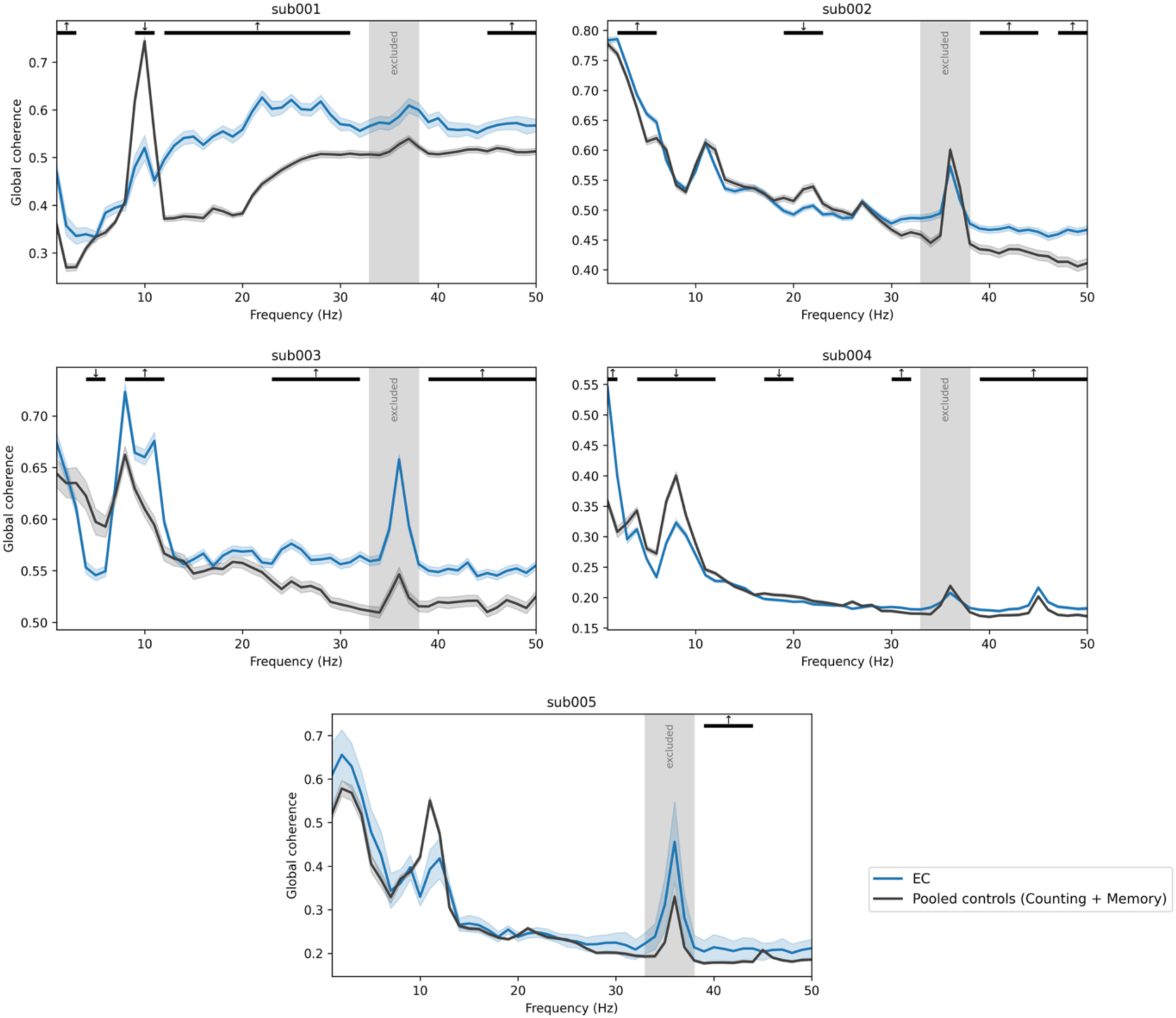
EC is associated with significant increases in high-frequency global coherence in all participants. Global coherence spectra (1–50 Hz, 1-Hz resolution) for each participant were computed as the ratio of the largest eigenvalue of the cross-spectral matrix to the sum of its eigenvalues at each frequency (28). Lines show the mean across non-overlapping 15-s epochs for the EC condition (blue) and the two control conditions pooled (black), shaded ribbons indicate ± 1 SEM across epochs, and epoch numbers per condition are given in **Table S1**. Black bars mark frequency ranges with significant differences between EC and control conditions under the cluster-based block-permutation test. Global coherence in the beta and/or gamma range was significantly higher on EC than the pooled controls for all participants. Increases in delta- to alpha-band global coherence also occurred in four participants. The gray band (33–38 Hz) was excluded from all inference: the 35–37 Hz notch filter applied during preprocessing renders coherence estimates in this range artifactual, producing the spurious coherence peak visible at 36 Hz in every participant. For sub002, the Counting and Memory conditions differed significantly from each other. Separate per-control comparisons are shown in **Figure S2**.

**Table 1.** Frequencies with significant changes in global coherence per subject. Direction indicates the sign of the difference (↑ EC > controls, ↓ EC < controls); ΔGC is the mean EC-minus-control global coherence across the cluster’s frequency bins; cluster mass is the summed −log_10_(p) across those bins; and the cluster p-value is corrected for multiple comparisons across frequencies by comparison against the maximum-cluster-mass distribution over permutations. *For sub002, the Counting and Memory conditions differed significantly from each other. Statistics for the separate per-control analysis are shown in **Table S2**.

| Subject | Direction | Cluster | $\Delta GC$ | Cluster mass | Cluster p |
| --- | --- | --- | --- | --- | --- |
| Sub001 | ↑ | 1–3 Hz | +0.087 | 11.1 | .0372 |
|  | ↓ | 9–11 Hz | −0.153 | 10.1 | .0436 |
|  | ↑ | 12–31 Hz | +0.135 | 69.1 | .0002 |
|  | ↑ | 45–50 Hz | +0.054 | 9.8 | .0454 |
| Sub002* | ↑ | 2–6 Hz | +0.028 | 12.7 | .0166 |
|  | ↓ | 19–23 Hz | −0.025 | 13.8 | .0148 |
|  | ↑ | 39–45 Hz | +0.036 | 12.8 | .0162 |
|  | ↑ | 47–50 Hz | +0.053 | 10.8 | .0228 |
| Sub003 | ↓ | 4–6 Hz | −0.055 | 7.6 | .0404 |
|  | ↑ | 8–12 Hz | +0.052 | 13.6 | .0052 |
|  | ↑ | 23–32 Hz | +0.036 | 34.4 | .0004 |
|  | ↑ | 39–50 Hz | +0.032 | 41.8 | .0002 |
| Sub004 | ↑ | 1–2 Hz | +0.140 | 7.4 | .0174 |
|  | ↓ | 4–12 Hz | −0.034 | 28.3 | .0002 |
|  | ↓ | 17–20 Hz | −0.009 | 8.7 | .0100 |
|  | ↑ | 30–32 Hz | +0.007 | 8.4 | .0114 |
|  | ↑ | 39–50 Hz | +0.011 | 39.0 | .0002 |
| Sub005 | ↑ | 39–44 Hz | +0.030 | 11.0 | .0198 |

Two participants exhibited decreases in beta-band global coherence: 19–23 Hz for sub002 and 17–20 Hz for sub005. Reductions in theta- to alpha-band global coherence occurred in three participants (9–11 Hz for sub001, 4–6 Hz for sub003, and 4–12 Hz for sub004) whereas increases in delta- to alpha-band global coherence occurred in four participants (1–3 Hz for sub001, 2–6 Hz for sub002, 8–12 Hz for sub003, 1–2 Hz for sub004).

We ran two control analyses to determine whether beta- and gamma-band global coherence are confounded by EMG artifacts. The first “spectral-extension” test confirmed a broadband, EMG-compatible power elevation during EC in three participants: relative to the pooled control conditions, power in the EMG-index band (62–95 Hz) of the raw sensor recordings was significantly elevated (all *p* < .0002) (**Figure S3**). In the same three participants, however, the cessation-related increase in the analysis band (39–50 Hz) significantly exceeded the broadband elevation (all *p* < .0002), demonstrating a genuine gamma-band component beyond what broadband EMG artifacts can produce (**Figure S3**). The remaining two participants showed no significant elevation in either the EMG-index band or the analysis band.

The second control analysis measured correlations between power in the analysis band during EC, divided by the pooled controls, and proximity to electrodes that are most susceptible to EMG artifacts. Four of the participants did not show a significant muscle-proximity gradient (ρ = 0.09– 0.41, spin-corrected *p* ≥ .11) (**Figure S4**). However, the gamma-band power of sub002 did correlate significantly with proximity to the EMG-maximal electrode sites (ρ = 0.45, spin-corrected *p* = .037). Altogether, these analyses indicate that tonic muscle activity generally contributes to, but does not account for, the high-frequency global coherence, except potentially in sub002.

### EC alters power inconsistently across participants

Changes in power within canonical frequency bands varied across participants. The most consistent effect was that beta and gamma power were significantly higher on EC in three participants: sub001, sub003, and sub004 (**Figure 3**; statistics are displayed in **Table 2**). EC was associated with significant decreases in alpha power in sub001 and increases in alpha power in sub003. Theta power was larger on EC in sub002, while delta power was larger for sub002 and sub004. For sub002, alpha, beta, and gamma power differed significantly between control conditions; per-control analyses revealed that delta and theta power for sub002 were greater on EC relative to Counting, but not relative to Memory (**Figure S5**). Sub005’s alpha power also exhibited significant differences between control conditions, but there were no frequency bands for which power on EC was significantly higher or lower than either control condition, or the pooled conditions. A supplementary analysis correcting power across frequencies rather than canonical bands is reported in **Table S2**.

**Figure 3.**
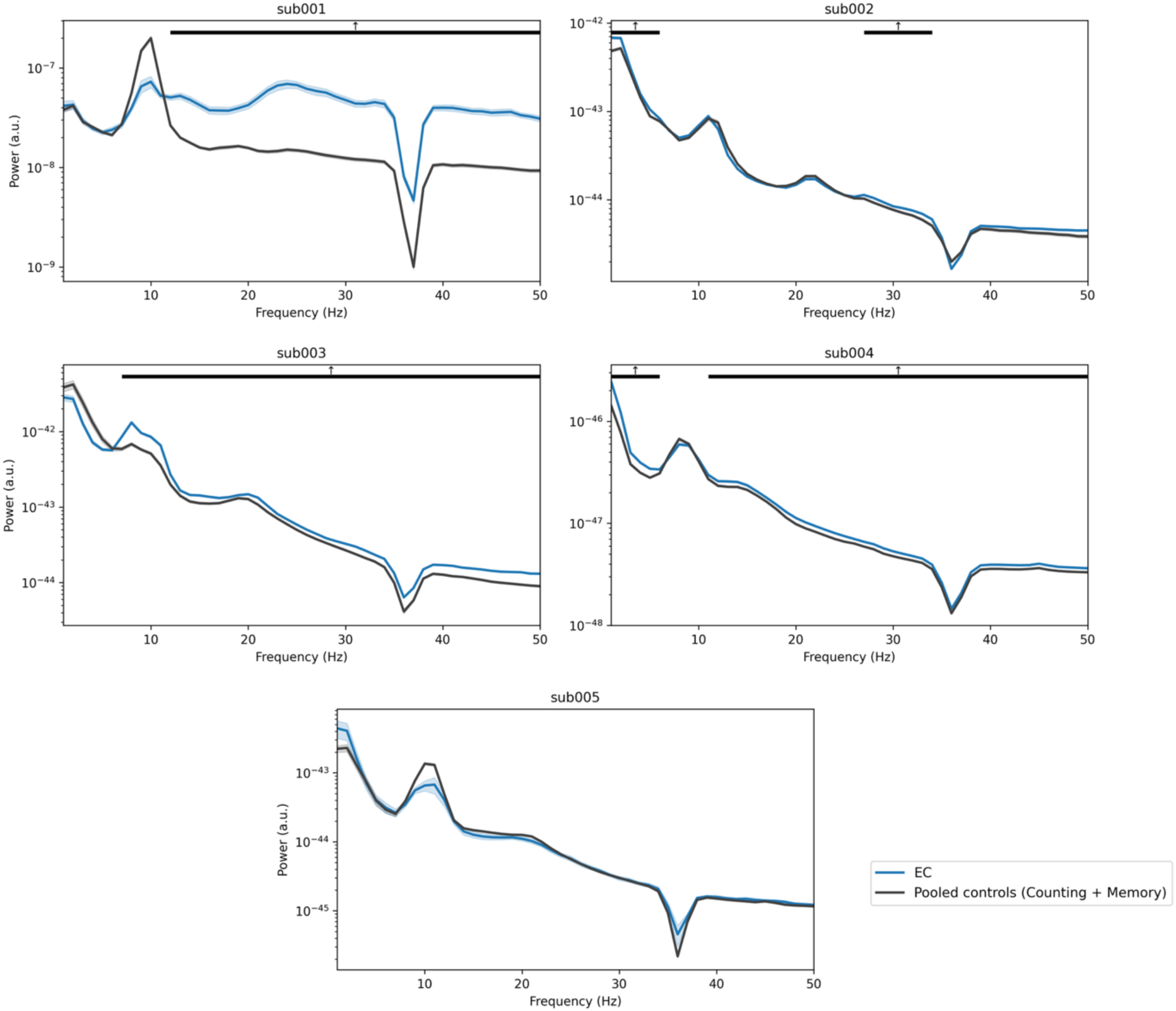
Changes in power vary across participants. Power spectra (EC, blue; pooled control conditions, black) were computed from the source-space timeseries of each participant. Shaded ribbons indicate ± 1 SEM across epochs; epoch numbers per condition are given in **Table S1**. Black bars mark frequency ranges with significant differences between EC and the pooled controls under the cluster-based block-permutation test. Beta- and gamma-band power was significantly higher on EC for three of the participants, whereas delta-band power was greater on EC for two participants. The narrow dip at 35–37 Hz visible in every spectrum reflects the notch filter applied during preprocessing. For sub002 and sub005, the Counting and Memory conditions differed significantly from each other for power measures; per-control comparisons are shown for all participants in **Figure S5**.

**Table 2.**
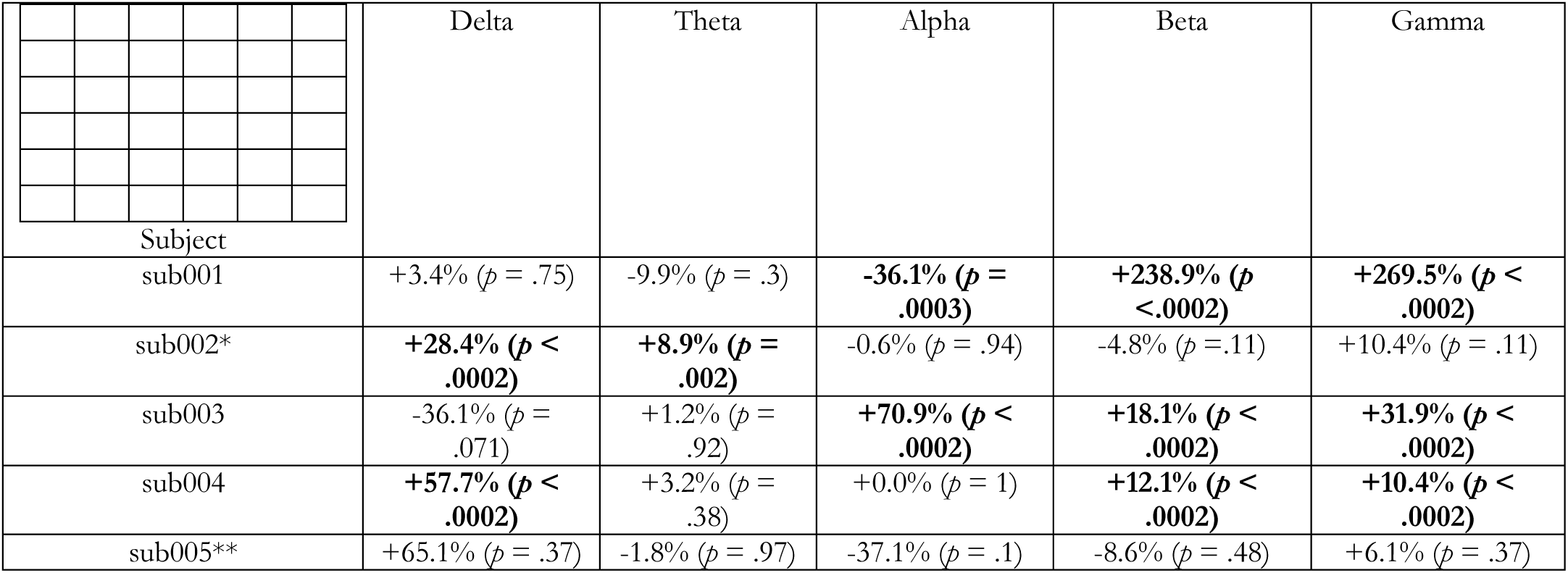
Changes in band power. Percentage change in band-averaged source power in the EC condition is shown relative to the mean of the two control conditions pooled, for the five canonical frequency bands (delta 1–4, theta 4–8, alpha 8–13, beta 13–30, gamma 30–48 Hz). p-values in parentheses are FDR-adjusted across the five bands within participant, with adjusted p < .05 in bold. *sub002’s control conditions differed significantly from each other in the beta and gamma bands. **sub005’s control conditions differed in the alpha band.

To localize the band-level effects, we measured the topography of gamma and beta power. The gamma-band increase was global in sub002-sub004 and more focal in sub005, where a cluster of right-parietal parcels increased against both control conditions (**Figure S6**). Beta-band increases followed a similar distribution (**Figure S7**). Across participants, the most consistent overlap lay in ventral somatomotor cortex, where beta and gamma power increased in all five participants. However, a regional version of the spectral-extension test showed that, in ventral somatomotor cortex, power also increased in the EMG-index band (62–95 Hz) in sub001, sub002, and sub003, identifying this region as the site of the broadband, muscle-compatible component rather than a neural signature. The topography of alpha power was also investigated due to prior literature about the role of alpha in cessation experiences. Left lateral inferior occipital alpha power decreased significantly in all participants except sub003 (**Figure S8**).

### EC does not consistently decrease complexity

EC was associated with significant increases in LZc relative to the pooled controls for sub001 and sub003, while sub004 exhibited a significant decrease (**Figure 4a**; statistics are displayed in **Table 3**). Sub002, sub003, and sub005 showed significant differences in LZc between control conditions. For sub002, LZc was significantly higher on EC than Counting only, and lower on EC than Memory; for sub003, LZc was significantly higher on EC than Memory only (**Figure S9a**). LZc was significantly greater on EC than both conditions (measured separately) and significantly lower on EC than both conditions (measured separately) for sub001 and sub004, respectively.

**Figure 4.**
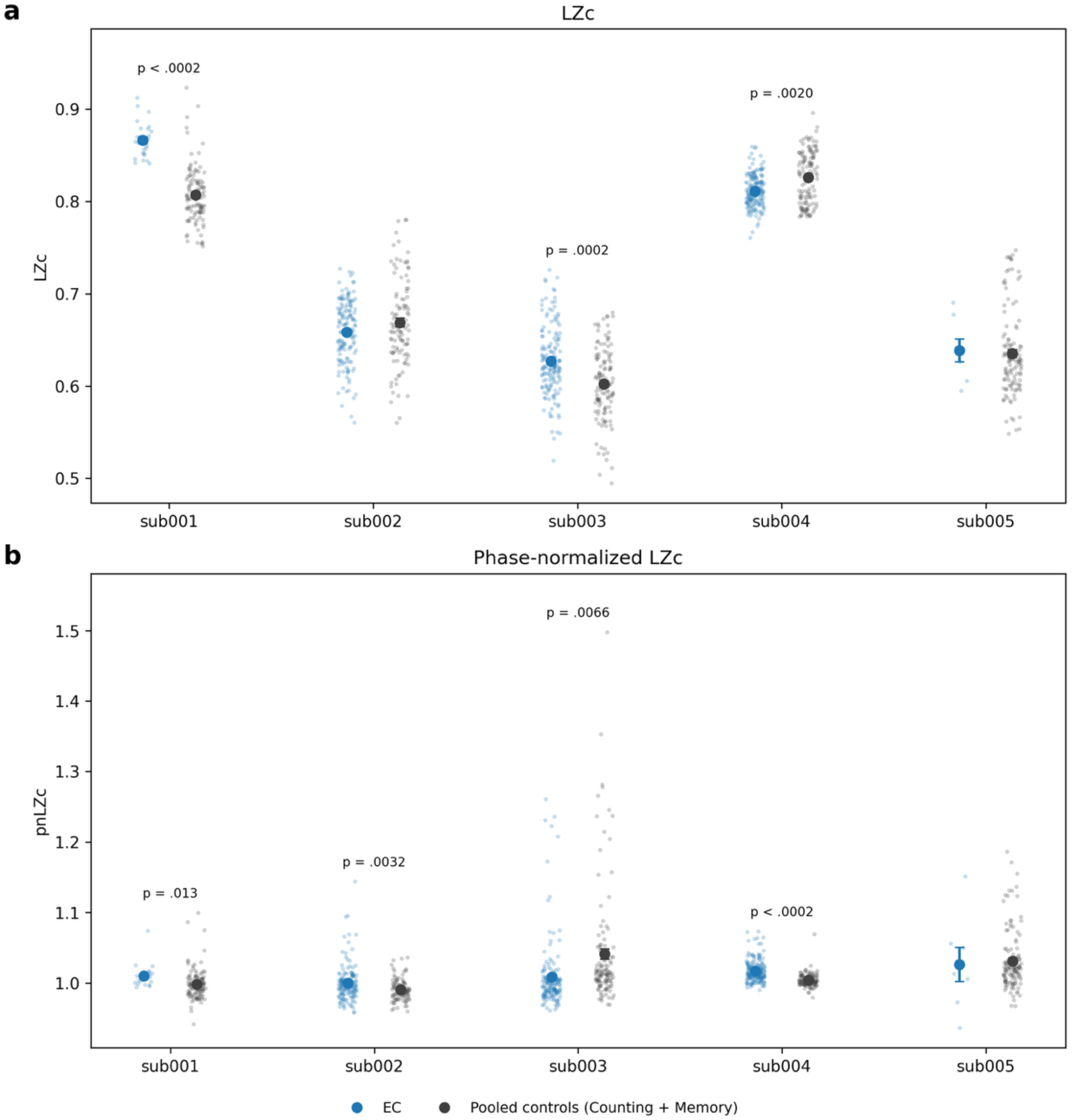
EC does not consistently reduce Lempel-Ziv complexity (LZc). (a) EC significantly reduced LZc relative to the pooled controls only in sub004, while significantly increasing LZc in sub001 and sub003. LZc significantly differed between conditions for sub002, sub003, and sub005; per-control analyses are shown in **Figure S9a.** (b) EC significantly increased phase-normalized LZc (pnLZc) relative to the pooled controls in sub001, sub002, and sub004, while decreasing it in sub003. pnLZc significantly differed between conditions for sub003; per-control analyses are showin in **Figure S9b**.

**Table 3.** Changes in complexity. Complexity was measured as Lempel-Ziv complexity (LZc), phase-normalized LZc (pnLzc), and Φ*, i.e., integrated information. Upwards arrows indicate significant increases on EC relative to the pooled conditions, and downward arrows indicate significant decreases on EC relative to the pooled conditions. † denotes that the Counting and Memory conditions differed significantly from each other for that participant and measure. Bolded p-values are significant; note that no correction was performed across the three

| Subject | LZc | pnLZc | $\Phi^*$ |
| --- | --- | --- | --- |
| sub001 | ↑ (**$p < .0002$**) | ↑ (**$p = .013$**) | ↑ (**$p < .0002$**) |
| sub002 | $p = .23 \dagger$ | ↑ (**$p = .004$**) | ↓ (**$p < .0002$**) |
| sub003 | ↑ (**$p = .0012$**) † | ↓ (**$p = .006$**) † | ↑ (**$p = .0010$**) † |
| sub004 | ↓ (**$p = .0018$**) | ↑ (**$p < .0002$**) | ↓ (**$p < .0002$**) |
| sub005 | $p = .93 \dagger$ | $p = .8$ | $p = .98$ |

Although changes in global LZc were not consistent across participants, there was a significant increase in LZc in left lateral inferior occipital cortex in all five participants during EC, relative to the pooled controls (**Figure 5**). This result held up in the per-control analysis, except for sub005, who only exhibited a significant increase relative to Memory. On the other hand, in four out of five participants, LZc decreased significantly in two regions within the default mode network, the left and right dorsomedial prefrontal cortex. One of these participants, sub003, only showed this decrease relative to Memory in the per-control analysis.

**Figure 5.**
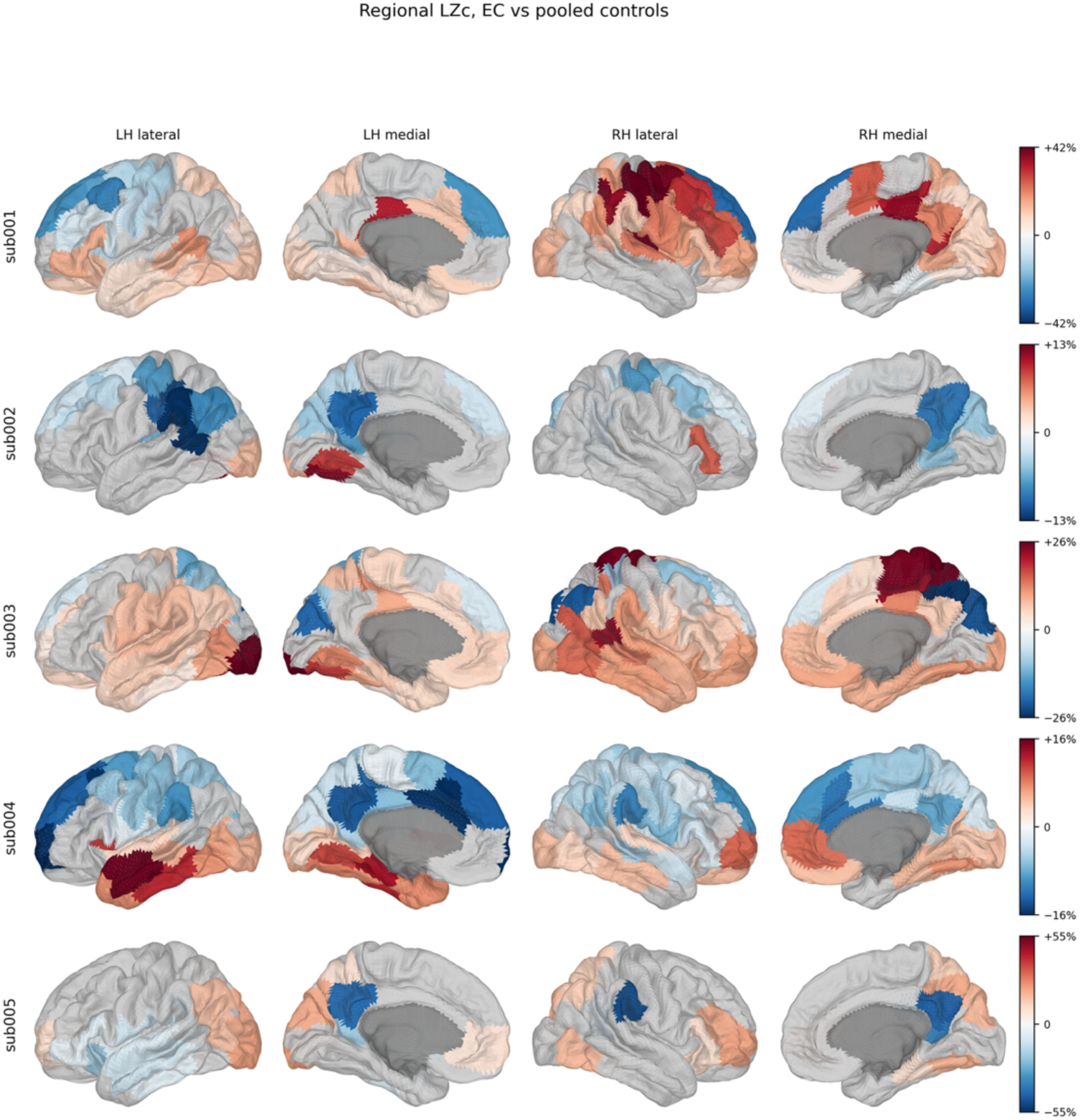
EC is associated with significant increases in regional Lempel-Ziv complexity in left lateral inferior occipital cortex. Percentage change in each parcel’s LZc in the EC condition is shown relative to the two control conditions pooled, displayed on the fsaverage5 surface (lateral and medial views of both hemispheres). p-values were corrected across the 100 parcels with the FDR; parcels not reaching adjusted p < .05 are shown in gray, and the medial wall in dark gray. Color scales are symmetric within participants. For sub002, sub003, and sub005, the Counting and Memory conditions differed significantly from each other in LZc. All five participants exhibit an increase in LZc in left ventral-occipital LZc, the same region that shows consistent alpha-power reductions (**Figure S8**), and four of five participants show decreases in LZc in frontal-midline default-network parcels.

Phase-normalized LZc (pnLZc) divides each segment’s LZc by that of a phase-randomized surrogate, which leaves the power spectrum identical but destroys all temporal structure beyond it. The ratio therefore indexes signal complexity that is not attributable to spectral content. Like their LZc, sub001’s pnLZc was significantly higher on EC than the pooled controls (**Figure 4b**; statistics are displayed in **Table 3**). However, the direction of the effect reversed for sub003 and sub004; pnLZc was significantly lower on EC for sub003 and higher for sub004. Meanwhile, sub002’s pnLZc was significantly greater on EC, whereas there was no significant difference in their LZc between EC and the pooled controls. In other words, the complexity changes were not reducible to spectral changes, but the relationship differed across participants. Sub003 exhibited significant differences in pnLZc between control conditions; their pnLZc was significantly lower on EC relative to Memory only (**Figure S9b**).

Finally, we also measured complexity with Φ*, which captures integrated information. Relative to pooled controls, Φ* was significantly greater during EC for sub001 and sub003, but significantly lower for sub002 and sub004 (**Figure S10**). Φ* and LZc display a consistent direction of effect in sub001, sub003, and sub004. There were no significant differences between control conditions in any of the participants; the per-control analysis is shown in **Figure S11**.

## Discussion

Here, we report the first evidence of neuroelectrophysiological correlates of self-reported extended cessation of consciousness in advanced meditators. Global coherence rose in the beta and/or gamma range during EC, with a significant increase somewhere between 39 and 50 Hz in all five meditators. Every other marker we examined behaved idiosyncratically. Band-limited power changed in different directions in different participants and in no case followed the canonical profile of pharmacological or physiological loss of consciousness. Neither Lempel-Ziv complexity (LZc), phase-normalized LZc (pnLZc), nor integrated information (Φ*) decreased consistently; the three measures frequently disagreed within the same participant. Regionally, however, two effects were highly consistent: LZc increased in left ventral/lateral inferior occipital cortex in all five participants, in the same territory that showed the most reliable alpha-power reduction, while LZc decreased in bilateral dorsomedial prefrontal cortex in four of five. Taken together, these data are difficult to reconcile with the view that EC leads to a global downregulation of cortical activity. They are more consistent with a targeted reorganization in which high-frequency activity becomes more globally coordinated, sensory cortex is not silenced, and complexity is redistributed rather than lost.

### EC increases high-frequency global coherence

Global coherence, as defined here, is the fraction of the cross-spectral matrix’s variance captured by its leading eigenvector at a given frequency (*28*). It is therefore a normalized measure of how strongly activity at that frequency is dominated by a single phase-locked spatial mode, not a measure of the absolute amount of connectivity. The finding that global coherence increased in the beta and/or gamma range in all five participants indicates that, during EC, high-frequency activity becomes more organized into a more dominant global mode.

This is not the signature that the anesthesia literature would predict. Propofol-induced unconsciousness and various disorders of consciousness are tracked most reliably by changes in *alpha-band* global coherence, which becomes strong and frontally dominant at loss of consciousness and collapses at recovery (*28*, *55*). In our data, low-frequency coherence was inconsistent across participants: increases in the delta-to-alpha range occurred in four participants and decreases in the theta-to-alpha range in three, with three participants (sub001, sub003, sub004) showing increases and decreases at different frequencies within this range. No participant showed the anesthesia-like pattern of a large, isolated alpha coherence increase. Intriguingly, however, ketamine-induced unresponsiveness is associated with a significant increase in high-beta and gamma-band global coherence (>27 Hz), though no control analyses were apparently performed to determine whether muscle artifacts contributed to this result (*56*). This raises the possibility that the EC shares a similar global coherence profile to anesthetic ketamine. Unlike EC, however, anesthetic ketamine can be associated with vivid hallucinations and dream-like states (*57*, *58*). Therefore, gamma-band global coherence may not index the presence of conscious content, but rather the decoupling of cortical dynamics from sensory input while large-scale high-frequency coordination is enhanced.

Long-range gamma-band synchronization is among the most frequently cited electrophysiological correlates of conscious access, whether framed as feature binding or as the late, widely distributed ignition posited by global workspace accounts (*59–61*). EC presents a unique case in which whole-brain gamma-band synchronization is enhanced while the participant reports that experience was absent altogether. Three interpretations are possible. The first is that large-scale high-frequency coordination is not sufficient for consciousness, and that these measures track the coordination of cortical activity rather than the presence of experience. The second is that the reported absence of experience is an absence of *recall* rather than of experience itself; because EC cannot be probed in real time, our phenomenological data are necessarily retrospective, and a state that is experienced but not encoded would be reported identically (*62*). The third is that the dominant high-frequency mode reflects a maintained but contentless configuration; in other words, high-frequency coordination remains stable but does not encode any conscious content. This last reading is speculative but is at least consistent with the participants’ descriptions of re-emergence: an objectless “pristine awareness” preceding the return of mind, body sensations, and inner speech.

Gamma-band EEG and MEG are vulnerable to EMG artifacts (*63*), but two analyses suggest that they do not completely account for the global coherence results. Firstly, although three of five participants did exhibit a significant increase of power in the 62–95 Hz EMG-index band during EC, this was exceeded by an increase of power in the 39–50 Hz analysis band. Since tonic EMG is spectrally broadband, a muscle artifact would have raised power in the analysis band and the EMG-index band to a similar degree (*43*). Secondly, the topography of power in the analysis band was not significantly aligned with the cranial musculature in all participants except sub002. The global coherence results for sub002 should therefore be treated with caution.

### EC does not reproduce the spectral profile of sleep or anesthesia

Loss of consciousness is typically associated with increases in low-frequency activity (*21–23*). Sleep and disorders of consciousness also diminish alpha power; in particular, diminished absolute alpha power is one of the most reliable biomarkers for distinguishing a more severe unresponsive wakefulness syndrome from less severe minimally conscious state (*64–66*). General anesthesia tends to “anteriorize” alpha power, that is, suppressing alpha power in occipital regions of the brain while elevating alpha power in frontal regions (*67–69*). A similar topography is seen in N2 sleep spindles (one stage above deep sleep), which are predominantly localized to frontal midline regions (*70*).

Changes in power during EC were inconsistent across participants. If anything, there was mixed evidence that the spectral profile of EC is similar to that of other states of unconsciousness. High-frequency power (in the beta and gamma bands) increased in three participants, which typically does not occur in other states of unconsciousness like GABAergic anesthesia and sleep (*71*, *72*). However, ketamine-induced anesthesia is associated with a characteristic gamma burst pattern in which gamma alternates with slow-delta oscillations (*73*). We did not observe a consistent increase in slow-wave power during EC, and our epoch-averaged analyses would not resolve such a temporal structure, so whether the high-frequency increase during EC is tonic or intermittent remains untested. Setting this aside, EC shares several features with anesthetic ketamine: elevated high-frequency power and increased high-frequency global coherence.

Meanwhile, during EC, alpha power at the band level changed significantly in only two participants and in opposite directions. The data therefore did not provide support for our hypothesis that EC would suppress alpha power, based on prior case reports of momentary cessations (*9*, *17*). Yet one of these case reports did localize pre-cessation alpha reduction to occipital cortex, which is consistent with our findings; four out of five participants exhibited a significant decrease in left lateral inferior occipital alpha power during EC. Reduced occipital alpha is a feature of most states of diminished consciousness: it is lost at induction of GABAergic anesthesia, under anesthetic ketamine, and in disorders of consciousness (*67–69*, *73*, *74*). However, under propofol and sevoflurane, occipital alpha loss is coupled to the emergence of a large, coherent frontal alpha oscillation (*67–69*), whereas under ketamine, it gives way to widespread coherent theta (*56*). Neither reorganization occurred in our participants; EC did not anteriorize alpha, nor did it globally increase theta power.

Altogether, EC’s spectral profile is neither that of GABAergic anesthesia nor a straightforward analogue of any other state. It shares the occipital alpha reduction common to most states of diminished consciousness, but not the frontal alpha that accompanies it under anesthesia. EC shares elevated high-frequency power and global coherence with anesthetic ketamine while lacking the slow-wave activity that they are typically associated with. EC appears to alter not the amount of activity in any band, but rather where and how that activity is organized; this theme recurs in the complexity findings below.

Automated sleep scoring further argues against the possibility that participants were simply asleep during EC (see **Supplementary Methods and Results**). Although delta power increased in two participants, slow waves were absent, and N2 sleep and spindles were no more frequent during EC than during the waking control tasks. REM sleep poses a subtler alternative: EC’s combination of reduced alpha and elevated beta and gamma activity resembles the activated EEG of REM sleep, and the classifier accordingly assigned a substantial fraction of sub004’s EC epochs to REM. Several considerations weigh against this interpretation. The classifier had access to EEG and EOG but not to the muscle atonia that defines REM sleep, and power in the EMG-index band increased rather than decreased during sub004’s EC; moreover, outside of narcolepsy, healthy adults do not transition directly from wakefulness into REM sleep. Finally, epochs scored as REM contained no more rapid eye movements than those scored as wake, and nearly all came from a single run with degraded electrode coverage over the vertex; tonic REM sleep, which lacks eye movements, cannot be excluded without chin EMG.

### Unlike other states exhibiting loss of consciousness, EC does not consistently alter neural complexity

According to the Entropic Brain Hypothesis, the richness of conscious content is encoded in the entropy, that is, the amount of variability or complexity, of brain activity (*75*). Psychedelics, which enhance the richness of conscious experience, increase the entropy of brain activity, though there have been difficulties replicating some of these findings in independent datasets (*48*, *76–79*). Depressed conscious states such as deep sleep, anesthesia, and disorders of consciousness, typically diminish LZc, permutation entropy, integrated information, and other indices of neural complexity, though there have been reports of increases in LZc after propofol-induced anesthesia (*24–27*, *80–82*).

One would therefore expect EC, which leads to an absence of consciousness, to decrease neural complexity. However, we did not find evidence for this hypothesis. LZc decreased significantly in only one participant while increasing significantly in two others. Meanwhile, pnLZc increased significantly in three participants while decreasing significantly in one. Thus, in one participant (sub003), increases in LZc were driven by changes in power, whereas in three other participants, EC altered the temporal structure of the signal beyond what its power spectrum would predict. Finally, Φ* increased significantly in two participants and decreased significantly in two others. Only sub001 showed a coherent direction across all three measures, and sub005 showed none. EC therefore does not reliably reduce neural complexity.

The lack of a consistent decrease in complexity may reflect the intensely concentrated state that is required to enter EC. While unconsciousness induced by sleep or anesthesia arises passively or exogenously, EC is voluntarily cultivated and typically preceded by advanced meditation training. In this study, all participants engaged in ACAM-J prior to entering EC, and most also incorporated elements of AIIM. These preparatory practices involve not only sustained attentional control, but also refined phenomenological insight into the mind. In general, focused attention meditation is an activity that requires meditators to consistently deviate from resting-state activity, since the mind wanders during the resting state and the meditation leads participants to return their concentration to the breath whenever they become distracted. Thus, focused attention meditation increases LZc relative to resting-state, especially for more advanced meditators, whereas mind-wandering during meditation, which leads to a decline in concentration, decreases LZc (*83–85*). Moreover, previous work from our group has shown that ACAM-J elevates LZc (*31*). It is possible that, even though there is reportedly no consciousness during EC, the concentrated state in ACAM-J persists into EC; indeed, advanced meditators may be so well-trained in focused attention that it can arise without their conscious awareness.

Furthermore, LZc may be less an index of the richness of experience and more a reflection of deviations from resting-state activity. For instance, broadband LZc increases when participants are engaged in tasks and even when participants ingest caffeine, even though consciousness is not necessarily becoming “richer” in either case (*86*, *87*). Because EC is not a conventional resting state and its preparation instead demands a high degree of concentration, LZc could plausibly increase even though the richness of conscious experience is decreased.

Thus, EC presents a unique scenario in which consciousness is absent without a concomitant decrease in neural complexity. This challenges the Entropic Brain Hypothesis and other theories that claim that complexity is a sufficient condition for consciousness. In other words, consciousness may require not just complexity per se, but specific network configurations of complex activity that were intentionally disrupted during EC. EC elevates complex activity in *local* brain networks, but these activities never coalesce into the unified, metastable patterns associated with conscious perception and cognition. For instance, EC increases LZc in specific regions of the brain, such as the left lateral inferior occipital cortex, but sometimes decreases it in other areas, such as left and right dorsomedial prefrontal cortex. On the other hand, deep sleep and disorders of consciousness tend to homogeneously decrease complexity across the whole brain (*26*, *88*).

Notably, the left lateral inferior occipital cortex is the same region in which alpha power most conssitently decreased. Two methodologically independent measures of EC therefore converge on visual cortex. Given the standard interpretation of occipital alpha as an index of cortical inhibition, the joint pattern of reduced alpha and elevated local complexity imply greater excitability of visual cortex during EC, in spite of an absence of consciousness (*89*, *90*). The lack of visual experience during EC therefore cannot be explained by loss of sensory activity; it must instead be mediated by a downstream neural process.

Indeed, EC was associated with decreases in LZc in bilateral dorsomedial prefrontal cortex in four of five participants. These parcels sit within the default mode network, whose activity is closely tied to self-referential and autobiographical processing (*91*, *92*). Reduced complexity in this network is consistent with the subjective reports of the meditators, who stated that autobiographical memory, ego-centered cognitive processing, and a boundary between self and environment were absent during EC. Attenuating internally-oriented cognition may contribute towards loss of conscioiusness in EC, but more evidence is needed to confirm this account.

In summary, EC presents a unique case in which complexity is preserved or enhanced locally, especially in sensory areas, while experience is reportedly absent. Meanwhile, global changes in complexity are inconsistent across participants, suggesting that the spatial distribution of complexity may be more relevant for encoding loss of consciousness during EC.

### Relation to recent findings on advanced meditation

In recent years, the “third wave” of meditation research has investigated the flourishing of human potential that can lead to and result from advanced meditation (*2*, *7*, *8*, *93*). Profound changes in consciousness arising from advanced meditation, referred to in the scientific literature as meditative endpoints, are described across contemplative traditions with terms such as “awakening,” “enlightenment,” or “salvation” (*94–99*). Practitioners reporting these endpoints describe lasting transformations in awareness and self-related processing. These investigations contribute to a broader scientific effort to understand emergent and altered states of consciousness, which are increasingly recognized as widespread tools for improving mental health (*100*, *101*).

Prior advanced meditation research has focused on ACAM-J, and to a lesser extent, AIIM, both of which served as preparatory practices for EC. Across a series of intensive case studies, ACAM-J has been associated with reduced broadband EEG power; increased EEG LZc; and enhanced fMRI activity in the visual cortex and many other cortical and subcortical regions (*31*, *33*, *34*, *36*). Additionally, ACAM-J increased global integration or connectivity, though one analysis found that this became less prominent in the later ACAM-J (*31*, *32*, *35*).

EC converges with these preparatory states on two points. It reduces occipital alpha power, consistent with the increased visual-cortex activity observed during AIIM (*37*). The increase in high-frequency global coherence in all five participants parallels the increased global integration described during ACAM-J (*31*, *32*, *35*).

However, unlike ACAM-J, EC did not increase LZc, nor did it reduce broadband power. Additionally, ACAM-J’s effects on global integration in the brain been reported predominantly in lower frequencies and in fMRI connectivity, whereas EC’s are concentrated above 30 Hz. One interpretation is that the transition into EC preserves the globally integrated regime established during ACAM-J but shifts it to a frequency range that does not support the alpha- and theta-mediated communication associated with conscious content. However, this remains speculative.

### Limitations

This study is the first of its kind and, as such, comes with important limitations. The sample size was small (*n =* 5) since there are very few meditators who can achieve EC; thus, these results should be considered preliminary and not necessarily generalizable to broader populations. That being said, our subject-level statistical approach, which is found in studies with small samples both in primates and humans, ensured that we did not make invalid group-level inferences (*28*, *51–54*).

Periods of EC were self-reported in participants and were not verified independently. Although all participants were advanced meditators, we cannot be certain that consciousness was completely absent during these self-reported states. There was also inherent uncertainty in determining the precise onset and offset of EC, since the timing of consciousness cessation was based on participants’ subjective reports and behavioral cues. Small timing offsets could dilute or obscure transient neural changes around the state transition.

The use of non-meditative cognitive tasks as control conditions may introduce some ambiguity in interpretation, as observed differences between EC and the controls may reflect the absence of task-related cognition rather than neural signatures specific to EC. Future studies may consider incorporating consciously mediated states (i.e., advanced meditation without cessation) to more precisely isolate the neural correlates of consciousness in this context.

Statistical power varied markedly across participants. Sub005 contributed only five usable EC epochs, and that participant’s null results across power and complexity should be read as uninformative rather than as evidence of no effect. Sub002’s control conditions differed significantly from each other on several measures, complicating interpretation of the pooled comparison for that participant.

Residual muscle contamination cannot be fully excluded for the global coherence findings, particularly in sub002, and no dedicated EMG channels were recorded.

Finally, it is worth noting that the advanced meditators in our sample may also exhibit different patterns of brain activity compared to the rest of the population during sleep and anesthesia. To truly contrast the neural signatures of EC with those of other states characterized by a loss of consciousness, future studies should perform direct *within-subjects* comparisons between brain activity during EC and brain activity during these other states in advanced meditators.

## Conclusion

Although meditators do not report consciousness during EC, the neural correlates of this state do not appear to resemble unconsciousness. The one effect present in every participant was an increase in beta- and gamma-band global coherence, but ketamine-induced anesthesia may be the only unconscious state that shares this neural signature. Spectral power did not follow the profile of sleep or anesthesia, complexity did not consistently decrease, and the most reliable regional effects moved in opposite directions — complexity rising in occipital cortex, alongside reduced occipital alpha, and falling in dorsomedial prefrontal cortex. EC therefore appears to involve a reorganization of cortical dynamics rather than a global suppression of them, with sensory cortex left in a state of heightened excitability while self-referential systems become less complex in their dynamics. If these preliminary findings hold in larger samples, they may challenge theories that global complexity and high-frequency brain activity are sufficient for consciousness.

## Acknowledgments

The authors would like to thank Fernando Rosas and Bobby Tromm for their guidance on the statistical approach and measurement of Lempel-Ziv complexity in this study.

## Author contributions

M.D.S. conceptualized and supervised the study, edited the manuscript, and acquired funding. K.S. preprocessed and source-reconstructed the data, conducted analyses, and wrote the first draft of the manuscript. R.M.P. W.F.Z.Y., and T.S. acquired the data and edited the manuscript. R.M.P. also assisted with data preprocessing.

## Data and materials availability

Pre-existing functions were used to perform preprocessing and source reconstruction and to compute global coherence, power, and Lempel-Ziv complexity. The human participant data reported in this study cannot be deposited in a public repository.

## Supplementary Materials

## Supplementary Methods and Results

### Sleep staging and sleep-event detection

To assess whether participants were asleep during EC, we applied automated sleep staging and sleep-event detection to the raw sensor-level EEG of all five participants, including sub004, whose EEG was otherwise not used in the source analyses, using YASA (v0.7) (*50*). For each participant, electrodes approximating C3, Cz, and C4 were identified from the digitized electrode positions: the electrode closest to the vertex (Cz) and the electrodes closest to 36° left and right of it along the coronal arc (C3, C4), all within 12° of the nominal positions. Data were re-referenced to the average of all channels with non-outlying variance, resampled to 100 Hz, and divided into non-overlapping 30-s epochs. Each epoch was staged with YASA’s pretrained classifier using C4 (C3 where C4 was unusable) and, where recorded, the electrooculogram. Because the classifier also uses the time elapsed since the start of the recording, which biases the beginning of any recording toward wakefulness, we held these time features constant across epochs; with them retained, the NREM results were unchanged and fewer epochs were scored as REM sleep. Sleep spindles (12-15 Hz) and slow waves (0.3-1.5 Hz; peak-to-peak amplitude 75-350 µV) were detected on C3, Cz, and C4 with YASA’s default criteria and expressed as events per channel per minute. Because the control conditions were waking tasks, their scores provide an empirical estimate of the classifier’s false-positive rate in these daytime recordings. EC was compared with the pooled controls using the block-permutation procedure described below, with 60-s blocks of two epochs. To evaluate epochs scored as REM sleep, we detected rapid eye movements (YASA, default criteria) on a horizontal electrooculographic derivation: the dedicated EOG channel most strongly correlated with the frontal EEG difference F8-F7 or, where no EOG channel tracked it (|*r*| < .5; all participants except sub003), F8-F7 itself, with F7 and F8 located geometrically as above. We also counted high-amplitude transients over the vertex (≥40 µV, 0.5-10 Hz), characteristic of drowsiness and N1 sleep, using the average of the usable electrodes surrounding Cz.

To ensure that participants were not simply asleep during EC, we measured the presence of slow waves, the defining feature of deep (N3) sleep. These were essentially absent in every participant during both EC and the control conditions (at most 0.2 per minute), and no EC epoch was scored as N3. Epochs scored as N2 sleep were rare during EC (0–9% of epochs in sub001–sub004; one of sub005’s six EC epochs) and were no more frequent than during the waking control tasks (0–8%; no participant showed a significant difference). Spindle detection rates during EC fell within the range observed during the control tasks (0–2 per minute, below the several per minute typical of N2 sleep), were significantly lower during EC than during the controls in sub002 (p = .013), and did not differ in sub003–sub005. The only significant increase occurred in sub001 (0.95 vs. 0.25 per minute, p = .002), none of whose EC epochs were scored as N2; this is most parsimoniously explained by the shift of this participant’s alpha activity toward 12–13 Hz, which overlaps the spindle detection band. However, the classifier assigned a substantial fraction of EC epochs to REM sleep in sub004 (30%, versus 1% during the controls; the fraction of epochs scored as any sleep stage was significantly greater during EC, p < .0002) and, non-significantly, in sub001 (29% versus 13%). These REM-scored epochs contained no more rapid eye movements than sub004’s EC epochs scored as wake (0.64 versus 0.67 per minute) and no vertex sharp waves, and 26 of the 28 came from a single run in which Cz, C4, and most electrodes surrounding the vertex were unusable, so that staging relied on C3 alone.

## Supplementary Figures and Tables

**Figure S1.**
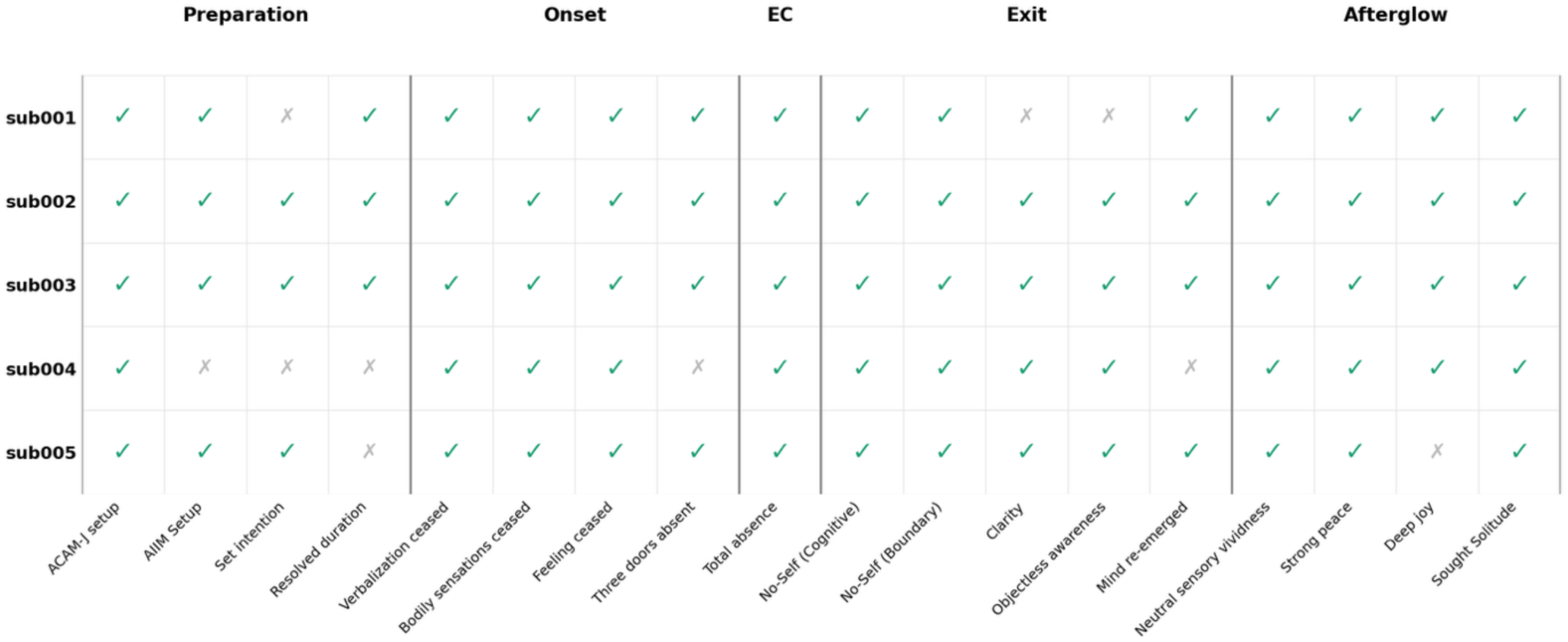
Subjective reports of different stages of EC are consistent across participants. Phenomenological items of EC were endorsed similarly across all participants. The categories in which participants differed were: whether advanced investigative insight meditation was done during preparation; a set intention to achieve EC; determination of duration of EC; absence of the perception of the three doors (i.e., incongruence, transient nature of experience, or self-lessness) prior to EC; experience of clarity and objectless awareness upon exiting EC, as well as re-emergence of the mind, bodily sensation, and internal verbalization; and experience of deep joy in the afterglow of EC.

**Figure S2.**
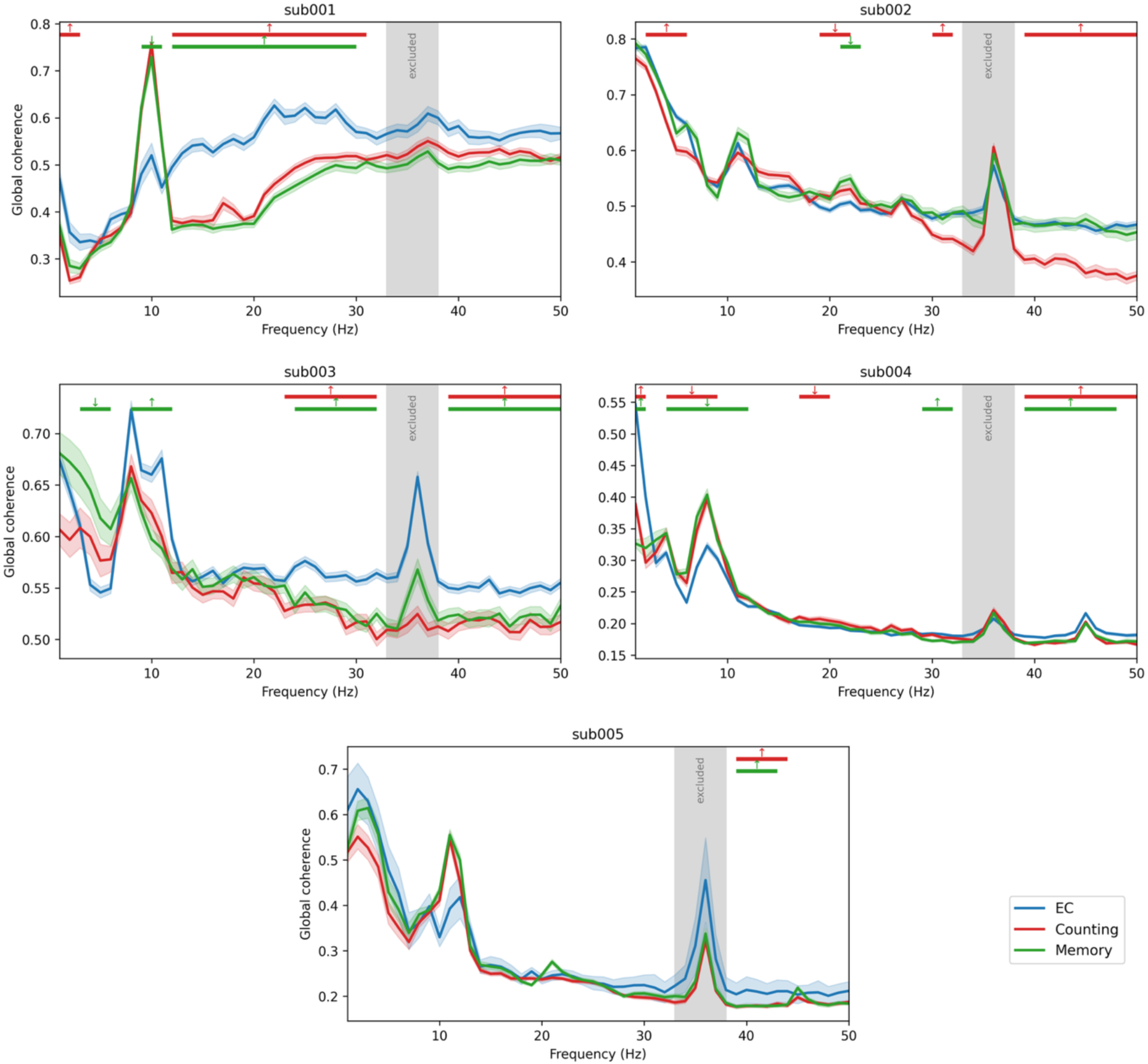
Per-control analysis of global coherence. Global coherence in the EC condition was compared to each control condition (Counting and Memory) separately. Red (green) bars mark frequency ranges with significant differences between EC and Counting (Memory) under the cluster-based block-permutation test. The gray band (33–38 Hz) was excluded from all inference: the 35–37 Hz notch filter applied during preprocessing renders coherence estimates in this range artifactual, producing the spurious coherence peak visible at 36 Hz in every participant. For sub002, the Counting and Memory conditions differed significantly from each other. Global coherence in the gamma band was significantly greater than only the Counting condition.

**Figure S3.**
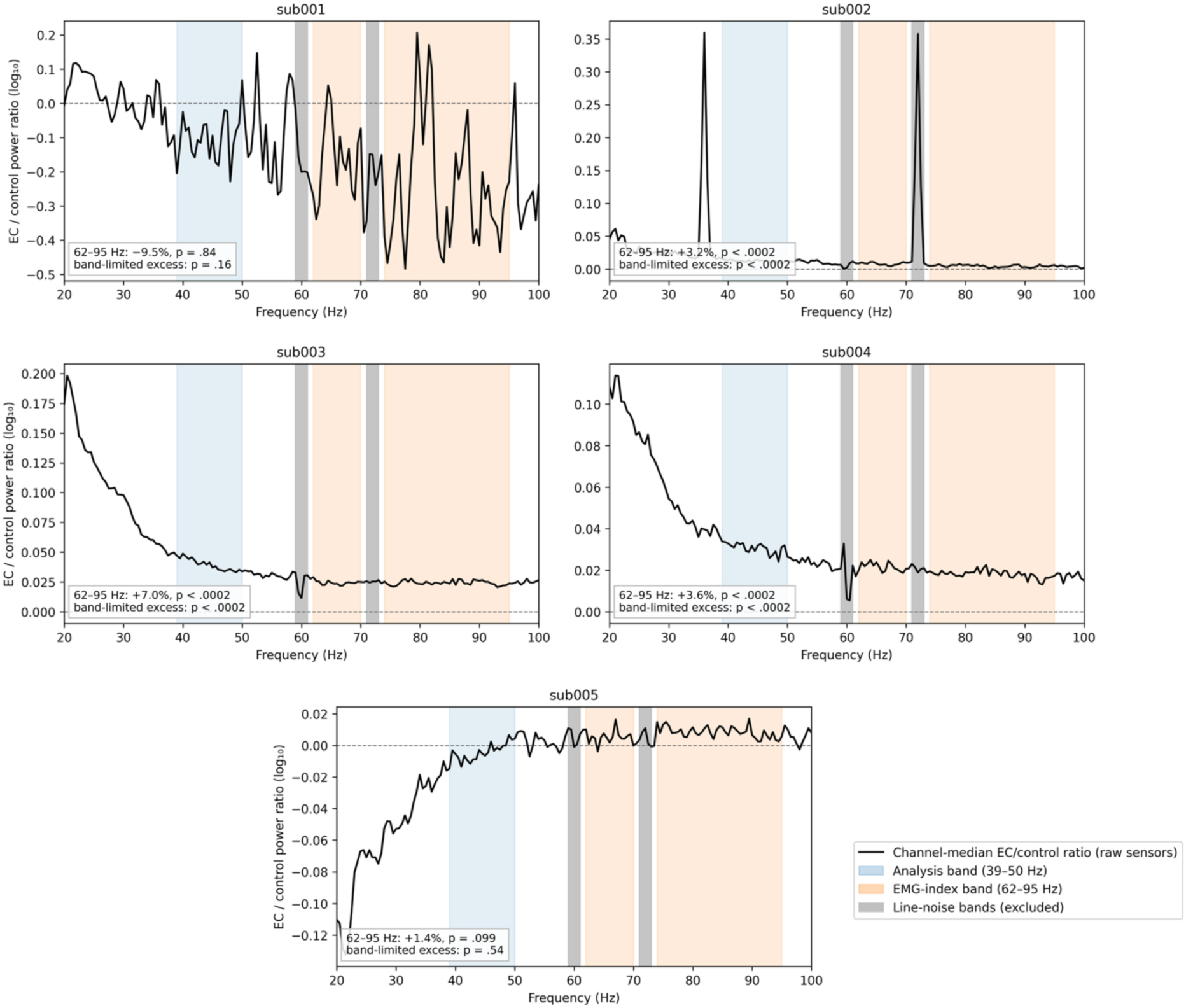
Spectral-extension tests for muscle contamination of high-frequency effects. We conducted two tests. First, we determined whether EC had significantly greater power than the pooled controls in an EMG-index band (62–95 Hz, orange). This test determines whether there is a broadband, EMG-compatible elevation in power. Second, we determined whether the ratio of EC to pooled-control power was greater in the analysis band (39–50 Hz, blue) than in the EMG-index band. If it is, then gamma-band activity should reflect genuine neural activity. The channel-median log_10_ ratio of EC to pooled-control power across 20–100 Hz was computed on the raw, unprocessed sensor recordings. Line-noise bands excluded from the EMG-index band (59–61 and 71–73 Hz, gray). A broadband elevation is present in sub002–sub004, indicating a partial myogenic contribution; in all three, the analysis-band increase nevertheless significantly exceeded it. The narrowband peaks in sub002 at 36 Hz and at its harmonic at 72 Hz (within the excluded line band) reflect that participant’s 33–39 Hz recording artifact. sub001’s power ratio spectrum is substantially noisier than the others’, reflecting uncleaned single-electrode EEG (versus MEG gradiometer medians) and its cross-session control recordings; both of its tests were null, with the control recordings carrying more high-frequency power than EC in the raw data.

**Figure S4.**
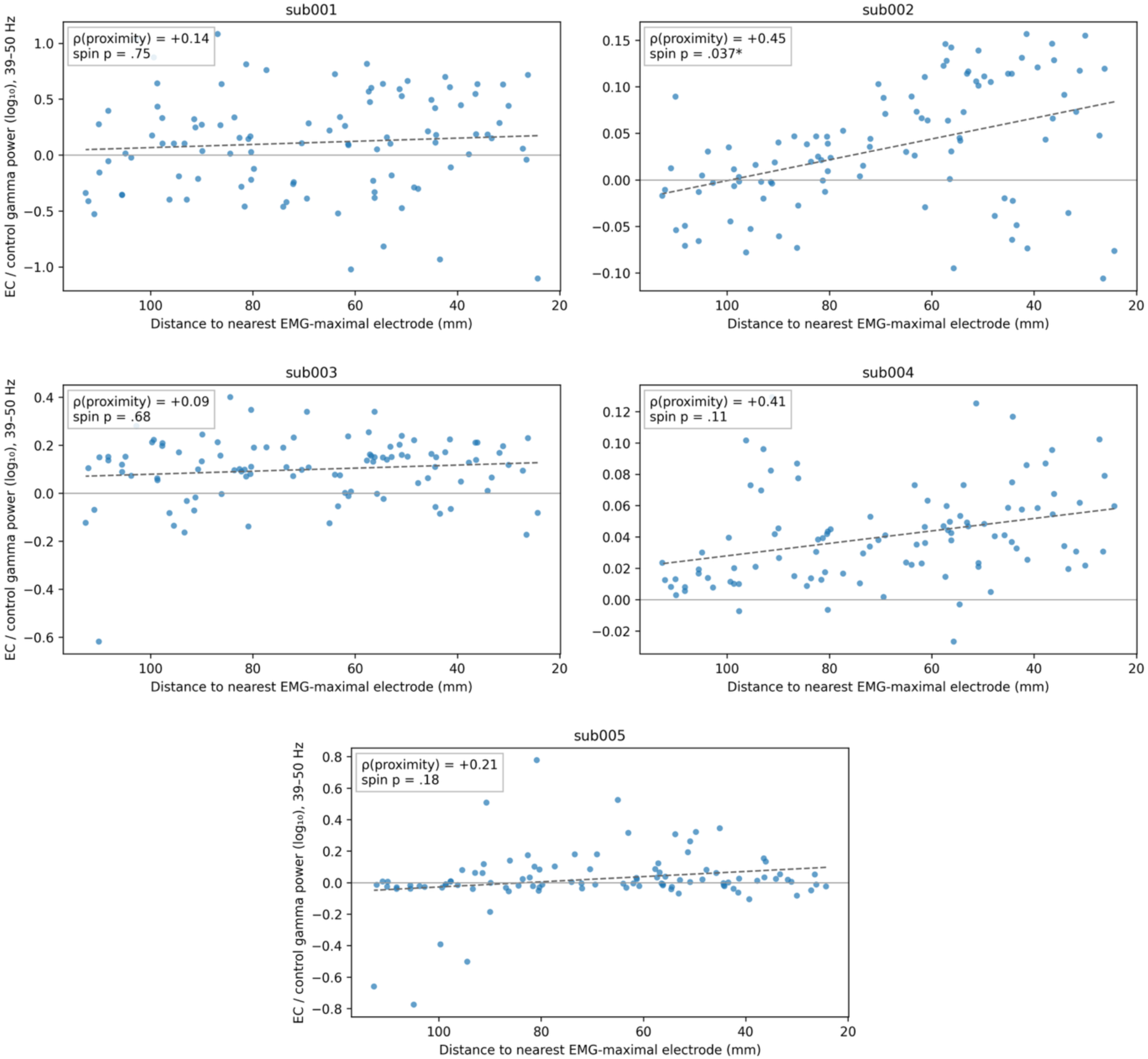
Spatial test for muscle contamination of high-frequency effects. Each point is one of the 100 Schaefer parcels; the vertical axis shows the parcel’s log_10_ EC/pooled-control gamma power ratio (39–50 Hz), and the horizontal axis its distance from the parcel’s MNI centroid to the nearest scalp site of maximal EMG contamination (electrodes Fp1/Fp2, overlying the frontalis, and F7/F8/T7/T8, overlying the temporalis). The x-axis is inverted so that muscle-proximal parcels appear to the right; dashed lines are least-squares fits. and ρ is the Spearman correlation between the gamma power ratio and muscle proximity. Significance was assessed with a volumetric spin permutation. Only sub002 showed a significant muscle-proximity gradient (*p < .05), converging with its other artifact indicators (**Figure S3**); no other participant approached significance, although gradients of |ρ| ≥ 0.30–0.57 were detectable under the rotational null. Correlations with radial (superficial-versus-deep) distance were null in all five participants (ρ = −0.22 to +0.17, spin p ≥ .19; not shown).

**Figure S5.**
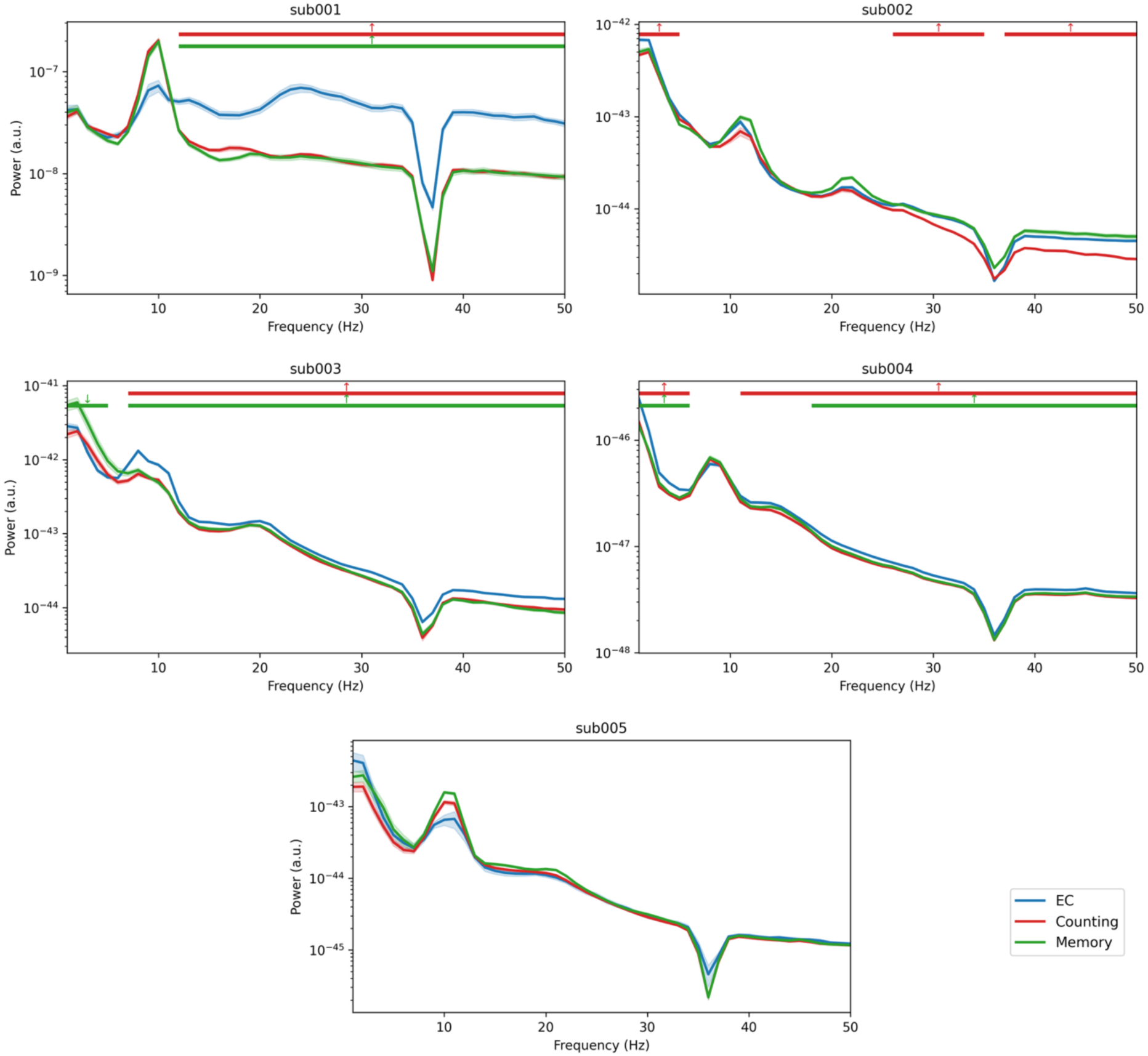
Per-control analysis of power spectra. Source power was compared between EC and each control condition separately. Shaded shaded ribbons indicate ± 1 SEM. Red bars mark frequency clusters with significant differences between EC and Counting, whereas green bars mark clusters with significant differences between EC and Memory. The narrow dip at 35–37 Hz reflects the band-stop filter applied during preprocessing and is present identically in all conditions. Delta and theta power for sub002 were greater on EC relative to Counting, but not relative to Memory.

**Figure S5.**
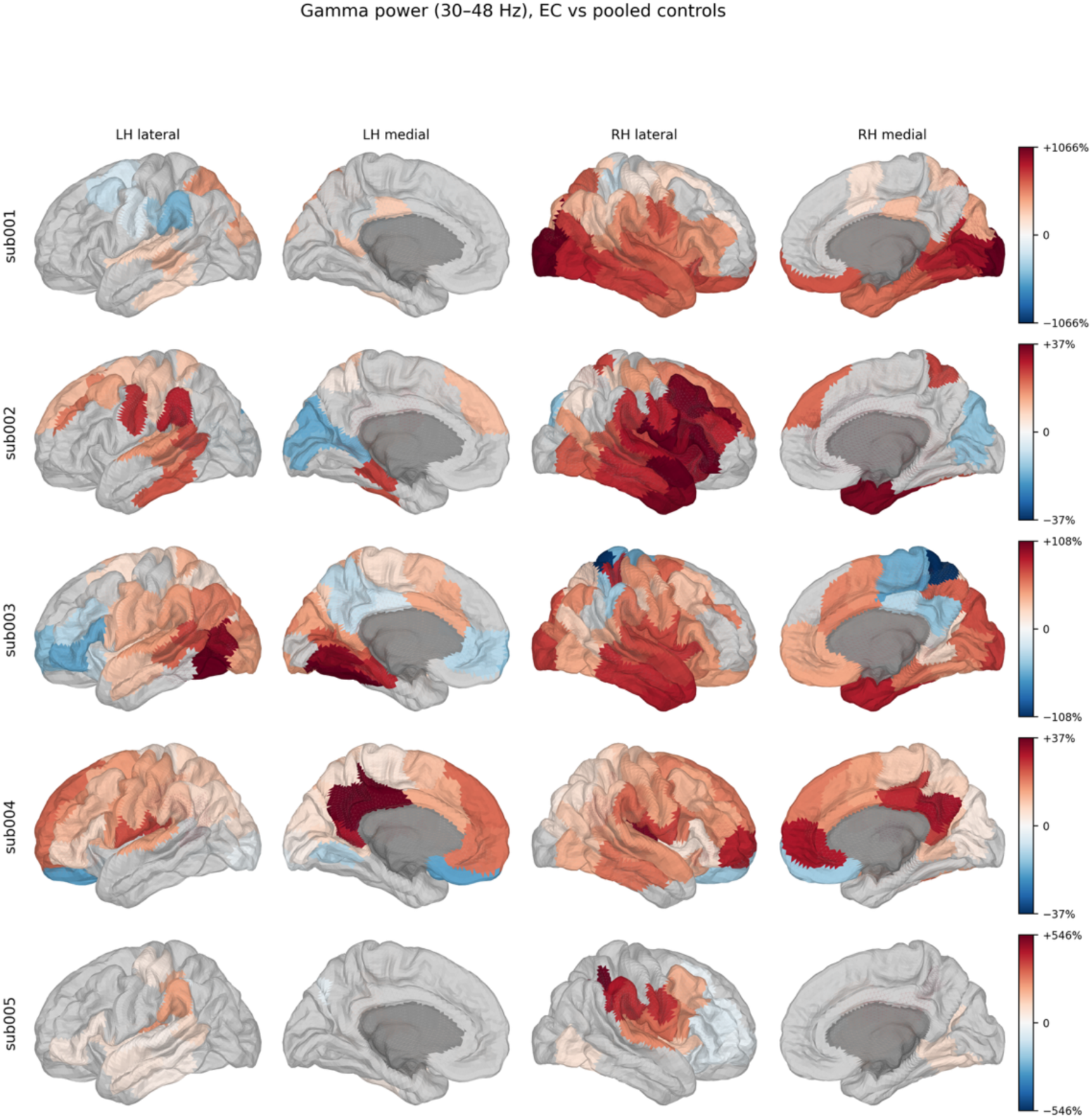
Regional gamma-band power changes during extended cessation. Percentage change in gamma-band (30–48 Hz) source power in the EC condition relative to the pooled control comparisons, per Schaefer-100 region, is displayed on the fsaverage5 surface. Parcel values were tested with block permutation and corrected across the 100 parcels with the FDR; parcels not reaching adjusted p < .05 are shown in gray, and the medial wall in dark gray. Note that, for sub002, regional gamma power differed significantly between control conditions. All participants exhibited increases in gamma power in ventral somatomotor cortex, but these increases extended into the 62–95 Hz EMG-index band and should not be interpreted as band-limited neural activity.

**Figure S7.**
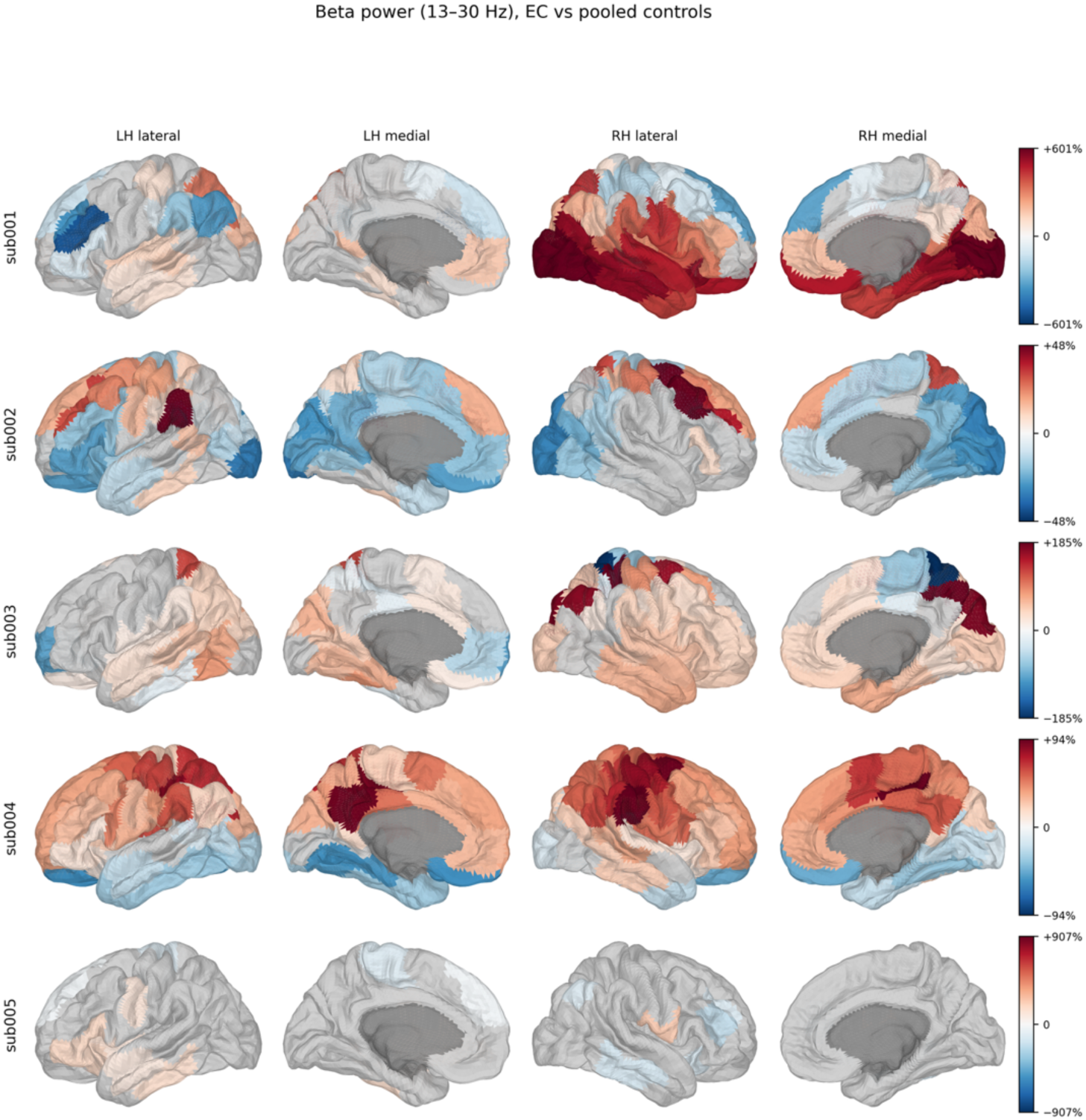
Regional beta-band power changes during extended cessation. Beta-band (13–30 Hz) regional source power is shown, measured with similar methods as **Figure S5**. Note that sub002 exhibited significant differences in regional beta power between control conditions. The same muscle-artifact caveat as in **Figure S5** applies to the ventral somatomotor increases, which were also observed in the regional beta power of all five participants.

**Figure S8.**
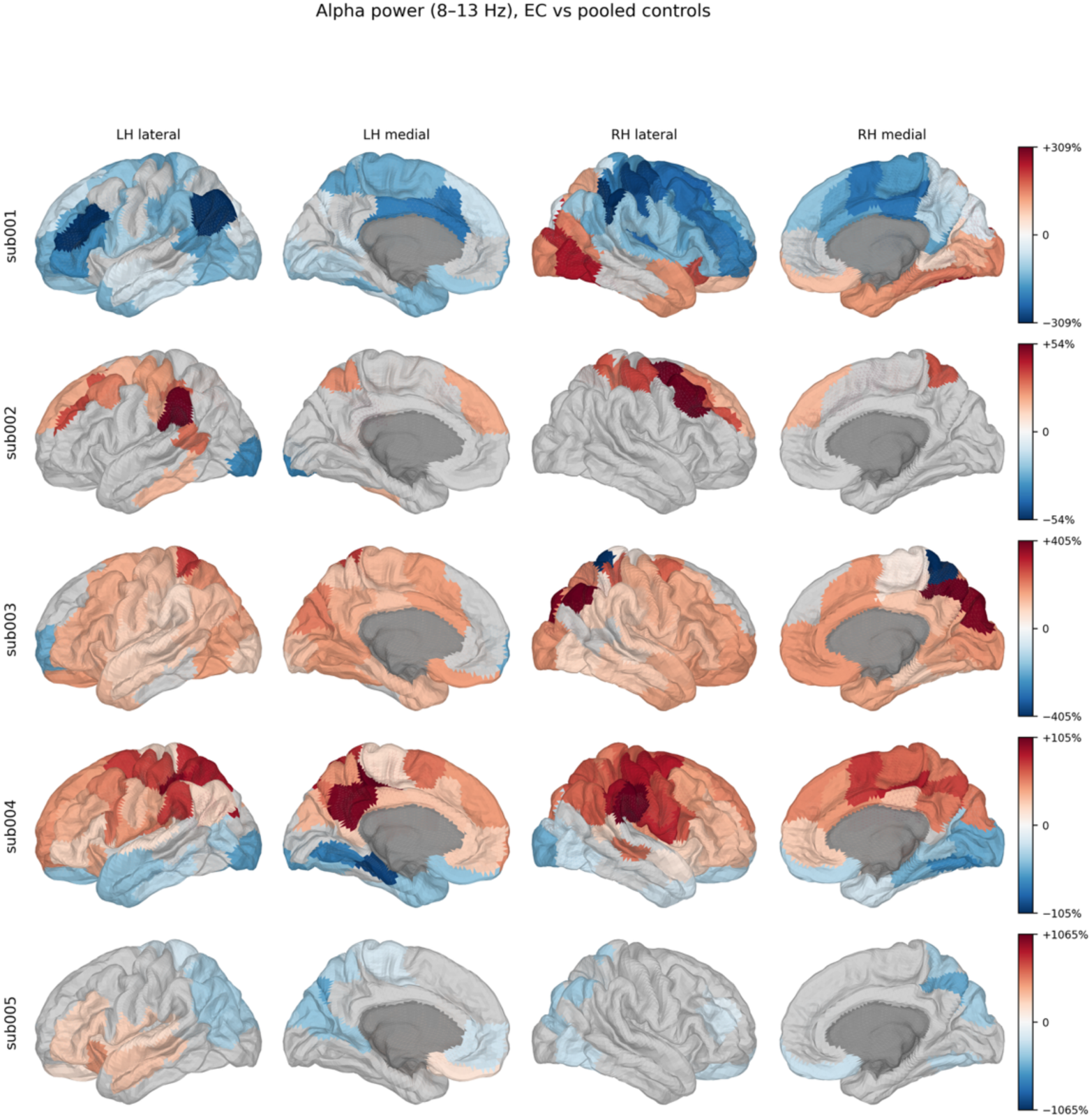
Regional alpha-band power changes during extended cessation. Alpha-band (8–13 Hz) regional source power is shown, measured with similar methods as described in **Figure S5**. Note that sub005 exhibited significant differences in regional alpha power between control conditions. Left ventral-occipital alpha reduction is shared by four of five participants.

**Figure S9.**
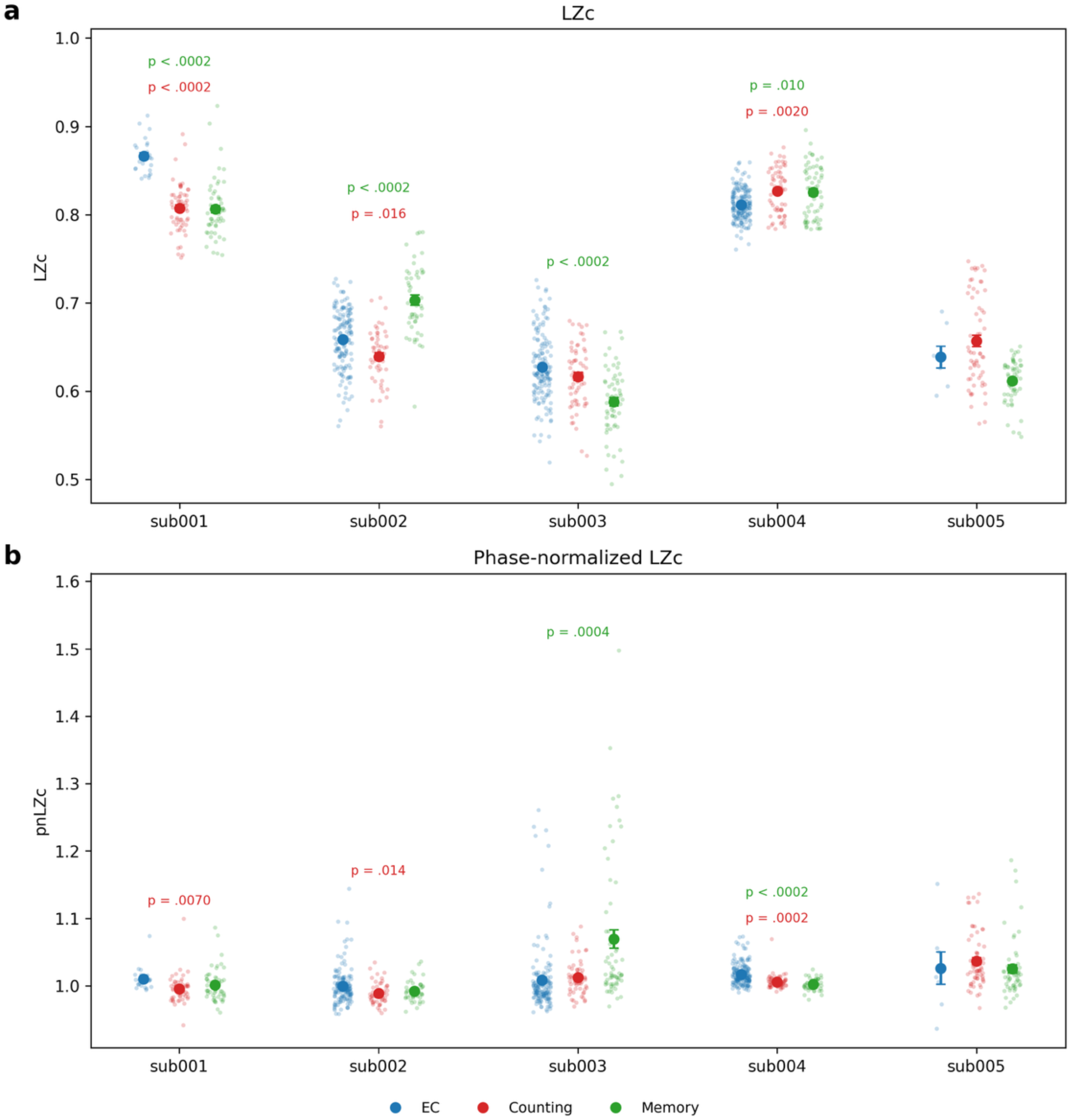
Per-control analysis of Lempel-Ziv complexity and phase-normalized Lempel-Ziv complexity. (a) Sub002, sub003, and sub005 exhibited significant differences in LZc between control conditions. During EC, sub001 exhibited significantly greater LZc relative to both control conditions; sub002 exhibited significantly greater LZc than Counting but significantly lower LZc than Memory; sub003 exhibited significantly greater LZc than only Memory; and sub004 significantly lower LZc than both control conditions. (b) Sub003 showed significant differences in phase-normalized LZc (pnLZc) between conditions. During EC, sub001 and sub002 each showed significantly greater pnLZc relative to Counting only; sub003 showed significantly lower pnLZc relative to Memory only; and sub004 showed significantly greater pnLZc relative to both control conditions.

**Figure S10.**
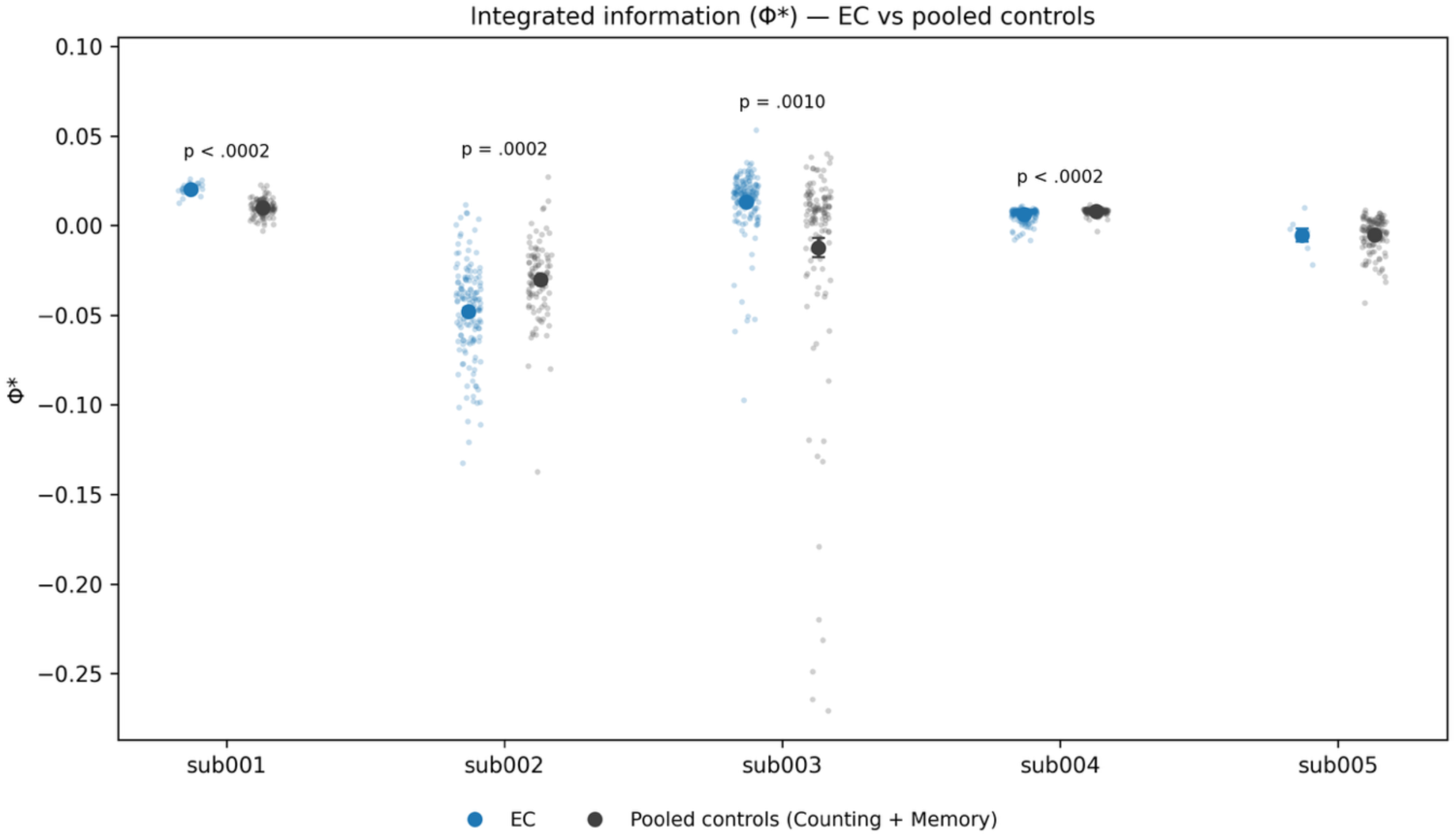
Changes in integrated information (Φ*) across participants relative to pooled controls. Integrated information was computed as Φ*: for every pair of the 100 parcels, we measured the time-lagged mutual information of the joint two-parcel system (lag of one sample, Gaussian assumption) minus the sum of the two parcels’ individual time-lagged mutual informations, averaged across all parcel pairs (49). Φ* quantifies the temporal information carried by the pair jointly beyond that carried by its parts and can take negative values when the parcels’ dynamics are redundancy-dominated.

**Figure S11.**
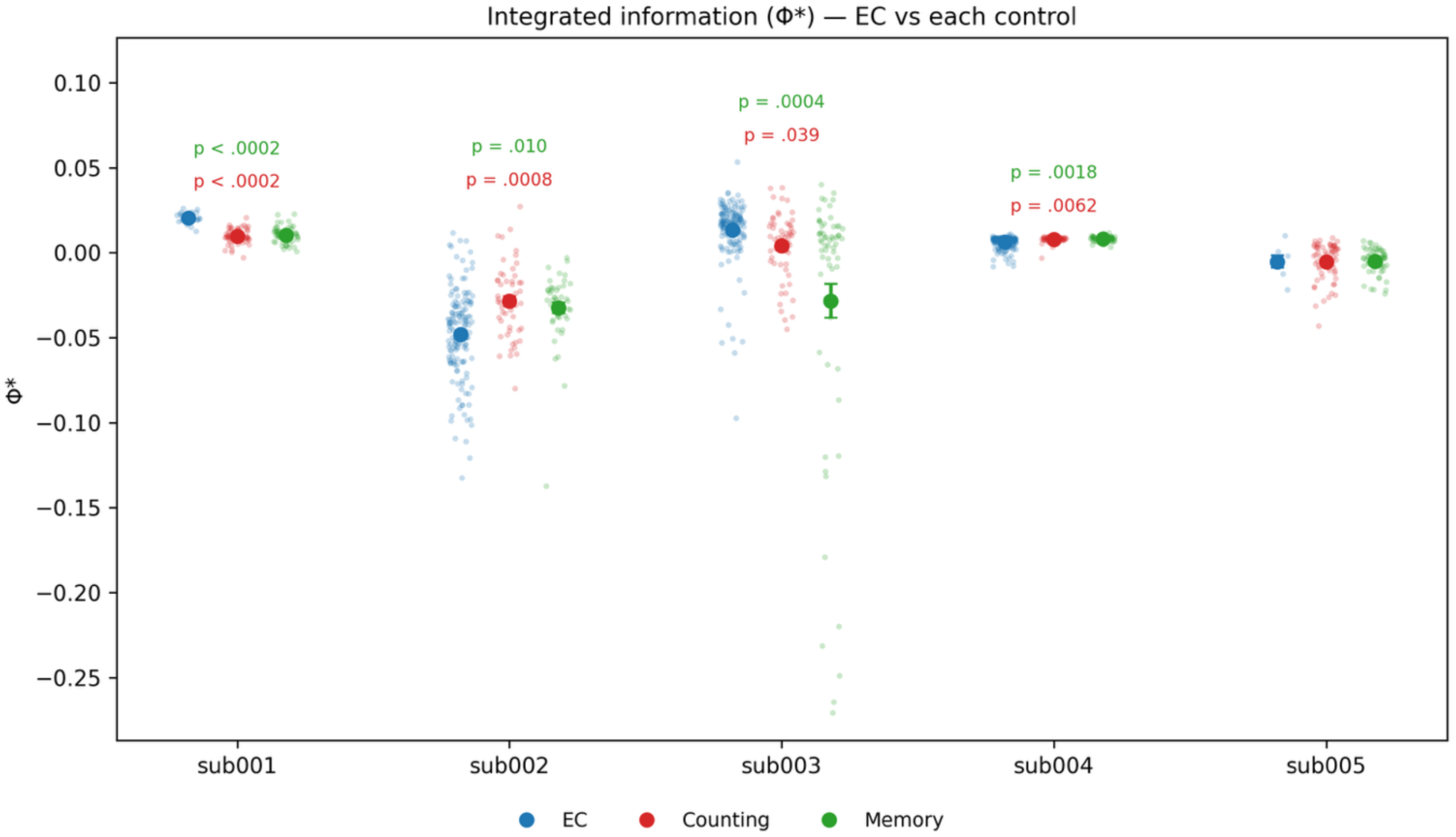
Per-control analysis of integrated information (Φ*). Integrated information was computed as Φ*: for every pair of the 100 parcels, we measured the time-lagged mutual information of the joint two-parcel system (lag of one sample, Gaussian assumption) minus the sum of the two parcels’ individual time-lagged mutual informations, averaged across all parcel pairs (49). Φ* quantifies the temporal information carried by the pair jointly beyond that carried by its parts and can take negative values when the parcels’ dynamics are redundancy-dominated.

---

**Table S1.** Run and epoch characteristics per participant. Number of runs of each condition, epoch length (which was always 15 seconds per epoch), and number of epochs created and epochs kept (after removing epochs with high variance)are shown.

| Subject | Run | Epoch Length (seconds) | Epochs created | Epochs kept |
| --- | --- | --- | --- | --- |
| sub001 | Counting 1 | 15 | 32 | 30 |
| sub001 | Counting 2 | 15 | 39 | 27 |
| sub001 | Memory 1 | 15 | 33 | 31 |
| sub001 | Memory 2 | 15 | 31 | 27 |
| sub001 | EC 1 | 15 | 27 | 26 |
| sub002 | Counting 1 | 15 | 59 | 56 |
| sub002 | Memory 1 | 15 | 58 | 48 |
| sub002 | EC 1 | 15 | 52 | 46 |
| sub002 | EC 2 | 15 | 61 | 51 |
| sub002 | EC 3 | 15 | 13 | 9 |
| sub002 | EC 4 | 15 | 51 | 42 |
| sub003 | Counting 1 | 15 | 32 | 32 |
| sub003 | Counting 2 | 15 | 31 | 29 |
| sub003 | Memory 1 | 15 | 32 | 31 |
| sub003 | Memory 2 | 15 | 31 | 31 |
| sub003 | EC 1 | 15 | 4 | 3 |
| sub003 | EC 2 | 15 | 67 | 65 |
| sub003 | EC 3 | 15 | 53 | 53 |
| sub003 | EC 4 | 15 | 33 | 31 |
| sub004 | Counting 1 | 15 | 32 | 31 |
| sub004 | Counting 2 | 15 | 33 | 32 |
| sub004 | Memory 1 | 15 | 31 | 30 |
| sub004 | Memory 2 | 15 | 32 | 31 |
| sub004 | EC 1 | 15 | 143 | 141 |
| sub004 | EC 2 | 15 | 34 | 33 |
| sub005 | Counting 1 | 15 | 33 | 32 |
| sub005 | Counting 2 | 15 | 33 | 26 |
| sub005 | Memory 1 | 15 | 32 | 31 |
| sub005 | Memory 2 | 15 | 32 | 31 |
| sub005 | EC 1 | 15 | 7 | 5 |
| sub005 | EC 2 | 15 | 3 | 2 |

**Table S2.** Changes in source power, corrected using cluster-based permutation testing across frequencies. This table shows all frequency clusters with a significant difference in source power between the EC condition and the two control conditions pooled (only clusters with corrected p < .05 are listed; sub005 showed none). Clusters were identified with the cluster-based block-permutation test over the full 1–50 Hz range. Direction indicates the sign of the difference (↑ means EC > controls); Δ power is the mean EC-minus-control power across the cluster’s frequency bins as a percentage of the control mean; cluster mass is the summed −log_10_(p) across the cluster’s bins. *sub002’s control conditions differed significantly from each other in the beta and gamma bands. **sub005’s control conditions differed in the alpha band.

| Subject | Direction | Cluster | $\Delta$ power | Cluster mass | Cluster p |
| --- | --- | --- | --- | --- | --- |
| sub001 | ↑ | 12–50 Hz | +241% | 144.3 | .0002 |
| sub002* | ↑ | 1–6 Hz | +27% | 18.1 | .0286 |
|  | ↑ | 27–34 Hz | +13% | 16.3 | .0340 |
| sub003 | ↑ | 7–50 Hz | +50% | 159.7 | .0002 |
| sub004 | ↑ | 1–6 Hz | +51% | 21.7 | .0298 |
|  | ↑ | 11–50 Hz | +11% | 143.3 | .0002 |
| sub005** | none | none |  |  |  |

